# Whole genome analysis of an extended pedigree with Prader–Willi Syndrome, hereditary hemochromatosis, and dysautonomia-like symptoms

**DOI:** 10.1101/019182

**Authors:** Han Fang, Yiyang Wu, Margaret Yoon, Laura T. Jiménez-Barrón, Jason A. O’Rawe, Gareth Highnam, David Mittelman, Gholson J. Lyon

**Author notes:** These authors equally contributed to this work (Han Fang, Yiyang Wu).

## Abstract

This report includes the discovery and analysis of a pedigree with Prader–Willi Syndrome (PWS), hereditary hemochromatosis (HH), and dysautonomia-like symptoms. Nine members of the family participated in whole genome sequencing (WGS), which enabled a wide scope of variant calling from single-nucleotide polymorphisms to copy number variations. First, a 5.5 Mb *de novo* deletion is identified in the chromosome region 15q11.2 to 15q13.1 in the boy with PWS. Second, a female invididual with HH is homozygous for the p.C282Y variant in *HFE*, a mutation known to be associated with HH. Her brother is homozygous for the same variant, although he has yet to be clinically diagnosed with HH. Third, none of the people with dysautonomia-like symptoms carry any reported or novel rare variants in *IKBKAP* that are implicated in familial dysautonomia (FD - HSAN III). Although two people with dysautonomia-like symptoms carry two heterozygous variants in *NTRK1*, a gene that has been shown to contribute to HSAN IV (congenital insensitivity to pain with anhidrosis, a disease that closely resembles FD), this variant is not present in the third proband. Fourth, WGS revealed pharmacogenetic variants influencing the metabolism of warfarin and simvastatin, which are being routinely prescribed to the proband. Finally, reports of the phenotypes were standardized with the Human Phenotype Ontology annotation, which may facilitate the search for other families with similar phenotypes. Due to the extreme heterogeneity and insufficient knowledge of human diseases, it is of crucial importance that both phenotypic data and genomic data are standardized and shared.

## INTRODUCTION

Many genetic tests have been commonly performed on individuals that have phenotypes overlapping with known diseases, especially for cancer and rare diseases (Meijers-Heijboer et al. 2000; Nanda et al. 2005; Sherman et al. 2005; Walker 2007). Physicians have also been routinely prescribing prenatal genetic tests and newborn screenings in clinics (Thompson et al. 2001; Morton and Nance 2006; Palomaki et al. 2011). However, there is a degree of uncertainty inherent in most genetic testing regarding the development, age of onset, and severity of disease (Evans et al. 2001). In addition, current genetic testing has not yet established predictive or even diagnostic value for common complex diseases (Smith et al. 2005). Some groups have begun to leverage the power of next-generation sequencing (NGS) to help diagnose rare diseases (Rope et al. 2011; Boycott et al. 2013; Koboldt et al. 2013; Honeyman et al. 2014). Many studies have used whole exome sequencing (WES) to facilitate the molecular diagnosis of individuals with diseases that appear to have a single large-effect size mutation contributing substantially to the development of the disease, referred to by some as “Mendelian disorders” (Bamshad et al. 2011; Lee et al. 2014). Of course, such disorders also have an extraordinary phenotypic variability and spectrum brought about by genetic background and environmental differences (Hamilton and Yu 2012; Li et al. 2012; Grillo et al. 2013; Lyon and O’Rawe 2014).

Despite much success using NGS-based techniques to identify mutations, there are still practical issues for the analytic validity for exome-or genome-wide NGS-based techiques, particularly in clinical settings (Lyon and Segal 2013; O’Rawe et al. 2013a). The clinical utility of genomic medicine is also uncertain, although some have suggested the need for better standards and benchmarking (Lyon 2012a; Dewey et al. 2014). Furthermore, research effort to date has been mostly driven by practicality and certain assumptions, such as focusing on coding regions, searching only for single-nucleotide polymorphisms (SNPs), or looking at a small set of known disease-relevant genes (Lyon and Wang 2012). However, the genetic architecture behind human disease is heterogeneous, and there are many reports of regulatory variants in the non-coding genome and splicing variants in the intronic regions that have a large-effect size on particular phenotypes (Slaugenhaupt et al. 2001; Faustino and Cooper 2003; Pagani et al. 2003; Venables 2004; Wang and Cooper 2007; Esteller 2011). In hypothesis-driven research studies, one might gain higher statistical power with a larger sample size by using cheaper NGS assays like WES or gene panels. But whole genome sequencing (WGS) has a unique strength in its ability to cover a broader spectrum of variants, small insertions and deletions (INDELs), structual variants (SVs), and copy number variants (CNVs) in studies where phenotype relevant variants might not be necessarily SNPs (Wang et al. 2013; Weischenfeldt et al. 2013; G. Day-Williams et al. 2015). In particular, WGS results in a more uniform coverage and better detection of INDELs, and is free of exome capture deficiency issues (Fang et al. 2014). When multiple human diseases segregate in the same family with distinct patterns, a more comprehensive genetic testing assay would be ideal, relative to targeted sequencing. Of course, cost and technical considerations have prohibited the wide adoption of highly accurate WGS for humans thus far, but this would indeed be the best assay to address the extreme heterogeneity of different genetic architectures for different diseases.

In line with other WGS efforts by the human genetics community, a report is given here of the discovery and comprehensive WGS analysis of an extended pedigree with Prader–Willi Syndrome (PWS), Hereditary Hemochromatosis (HH), dysautonomia-like symptoms, Tourette Syndrome (TS) and other illnesses.

## RESULTS

### Clinical presentation (with HPO annotation) and family history

Here, we present the phenotypic characterization of a Utah pedigree K10031, consisting of 14 individuals from three generations (Figure 1) with various medical conditions as mentioned above. The two probands we discuss in detail below come from two nuclear families in this extended pedigree.

**Figure 1.**
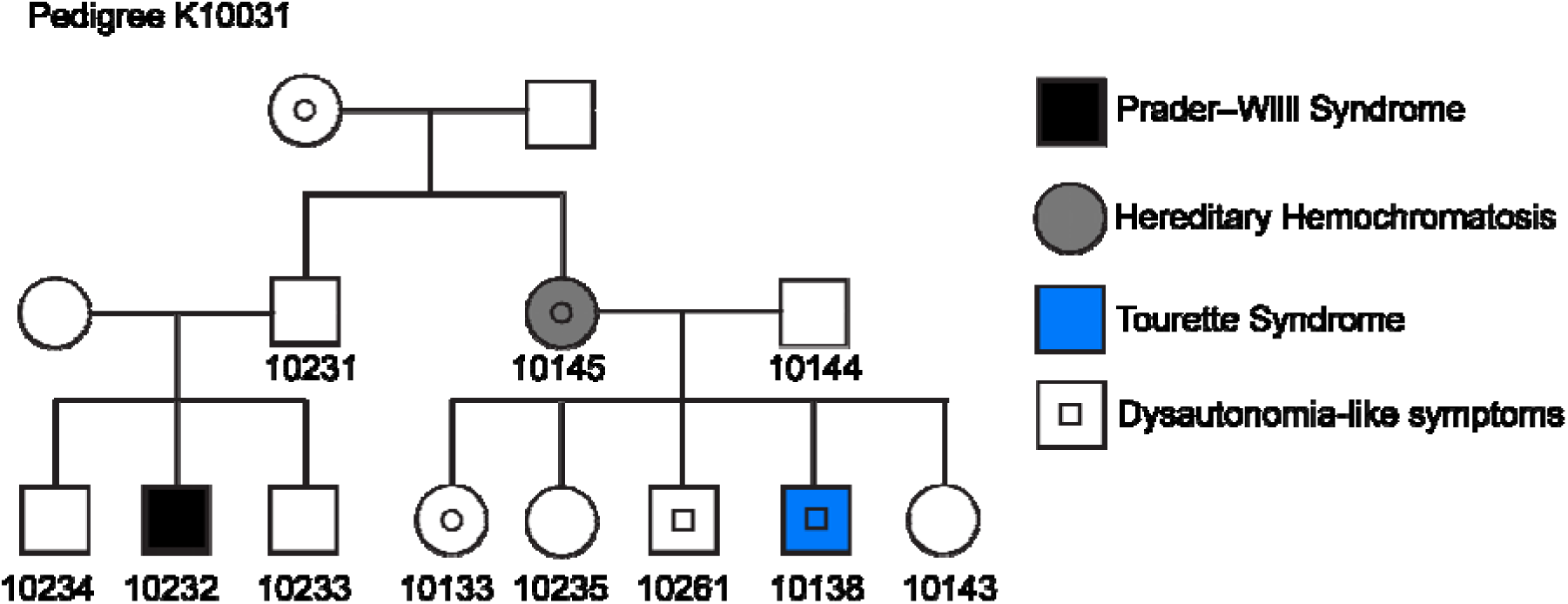
A pedigree spanning three generations with multiple rare diseases in this study. DNA was collected with informed consent from individuals marked with a number underneath underwent. All samples except K10031-10234 and K10031-10261 underwent WGS.

### The first proband K10031-10232

This proband (K10031-10232) is a 25-year-old (25 y.o.) male. He is the son of a Caucasian male, K10031-10231, and a Korean female. The mother did not participate in the study. He has two older male siblings, namely K10031-10233 and K10031-10234. The proband was diagnosed with PWS at 11 months old, and has dysmorphic facial features including a narrow forehead, downslanted palpebral fissures and almond-shaped eyes. A video recording (HDV_0073) of K10031-10232 explaining his medical presentation is described in the supplemental videos section and can be provided on request to qualified investigators. Since the PWS diagnosis, his behaviors have been assessed in great detail (Table 1, and Supplemental notes), from which the following diagnoses have been given: obsessive-compulsive disorder (OCD), depression, anxiety disorder, pervasive developmental disorder (PDD), hyperphagia, trichotillomania, and daytime hypersomnolence. He has an IQ ranging between 60-65, which is associated with mild mental retardation. He also has diagnoses of mild dysarthria, obstructive sleep apnea syndrome (OSAS), and severe scoliosis, and the latter has been corrected surgically. He has also undergone orchiopexy, tonsillectomy, and adenoidectomy. His physical exam is otherwise unremarkable. He has also denied having significant psychotic symptoms, including auditory or visual hallucinations, delusions, ideas of grandiosity, or paranoid ideation.

**Table 1.** Clinical Presentation of Proband K10031-10232

| **Table 1. Clinical Presentation of Proband K10031-10232** |  |
| --- | --- |
| **General Information** |  |
| Age (years) | 25 |
| Gender | Male |
| **Clinical Manifestations** | **HPO#** |
| **Prenatal History** |  |
| Cesarean section | 0011410 |
| Gestational diabetes | 0009800 |
| Oligohydramnios | 0001562 |
| Premature birth (36 weeks gestation) | 0001622 |
| **Development and Growth** |  |
| Delayed speech and language development | 0000750 |
| Dysarthria | 0001260 |
| Growth hormone deficiency | 0000824 |
| Poor fine motor coordination | 0007010 |
| Mild intellectual disability | 0001256 |
| **Facial Features** |  |
| Abnormal facial shape | 0001999 |
| Almond-shaped eyes | 0007874 |
| Downslanted palpebral fissures | 0000494 |
| Narrow forehead | 0000341 |
| **Other Physical Features** |  |
| Cryptorchidism | 0000028 |
| Excessive daytime sleepiness | 0002189 |
| Infantile muscular hypotonia | 0008947 |
| Obstructive sleep apnea syndrome | 0002870 |
| Scoliosis | 0002650 |
| **Behavior Features** |  |
| Aggressive behavior | 0000718 |
| Anxiety | 0000739 |
| Depression | 0000716 |
| Flipping pages in books | NL |
| Hair-pulling | 0012167 |
| Hyperesthesia | 0100963 |
| Impaired ability to form peer relationships | 0000728 |
| Impaired social reciprocity | 0012760 |
| Inflexible adherence to routines or rituals | 0000732 |
| Irritability | 0000737 |
| Lack of insight | 0000757 |
| Lack of peer relationships | 0002332 |
| Low frustration tolerance | 0000744 |
| Nail-biting | 0012170 |
| Obsessive-compulsive disorder | 0000722 |
| Pain insensitivity | 0007021 |
| Polyphagia | 0002591 |
| Poor eye contact | 0000817 |
| Restrictive behavior | 0000723 |
| Short attention span | 0000736 |
| Skin-picking | 0012166 |

**Abbreviations:** BMI- Body mass index, NL- Not listed.

Significant environmental risk was incurred when K10031-10232 was born at 36 weeks gestation through an emergency cesarean section (C-section) due to maternal failure to pass the nonstress test (NST). The pregnancy was significant for severe gestational diabetes and oligohydramnios, the latter serving as the original indication for the NST. His neonatal course was without complication, although some degree of neonatal hypotonia was noted. In regards to his family history, his paternal grandmother is reported to have anxiety disorder, and his father to have thrown vehement tantrums as a child. However the tantrums subsided during his developmental maturation.

At K10031-10232’s most recent clinical visit at 24 y.o., his height, weight, and BMI were 1.778 m, 79.83 kg, and 25.25 kg/m^2^ respectively. He appeared happy and was performing well on daily doses of Abilify and Zoloft for treating his OCD, and subcutaneously injection of Somatropin (gonadotropin) 6 days per week for his PWS. In adolescents and adults with PWS, low baseline and GnRH-stimulated gonadotropin levels are a frequently reported finding, arguing for an intrinsic hypogonadotropic hypogonadism. Most people with PWS receive this treatment. He has been using the continuous positive airway pressure (CPAP) machine during the past 6 months for his OSAS, and has been tolerating it well. His hyperphagia is continuously managed by parental education and structured, consistent parenting techniques. He is also exercising regularly by walking approximately six miles daily six days per week. Overall, from a PWS standpoint, he is functioning well and has transitioned successfully into a community setting, where he is employed at a laundry facility.

In an effort to help standardize phenotype reporting, we used Human Phenotype Ontology (HPO) annotation (Kohler et al. 2014). See Table 1 for a list of clinical phenotype features collected from the proband K10031-10232 with HPO annotations. For K10031-10232, the Phenomizer tool (Kohler et al. 2009) ranked the diagnosis for Prader-Willi Syndrome as the highest priority diagnosis (see Supplemental Data File 6 for the full report), supporting the fact that highly specific and annotated phenotype information can yield accurate diagnoses, at least for a clear syndrome like PWS. As presented below, the genomic analysis of K10031-10232 further confirmed deletions in the chromosome regions from 15q11.2 to 15q13.1, making PWS the most credible diagnoses for K10031-10232 at present.

### The second proband: K10031-10133

Proband K10031-10133 is a 26 y.o. female, born to a Caucasian mother (K10031-10145) and Caucasian father (K10031-10144). She is the eldest child amongst her two sisters and two brothers. Prior to age 18, K10031-10133 had a fairly unremarkable medical history. Arthralgia and episodes of fatigue and dizziness started at around 18 years of age. At age 20, she started to have refractory syncopal events, which led to multiple body injuries, and her cardiologist implanted a dual-chamber pacemaker (PM), after conducting standard workups. K10031-10133 was 21 years old at the time of the PM implantation. However, despite the PM placement, she continued to experience syncopal events, with a frequency of one episode a week starting at age 23. During the same time, in addition to the syncope, she also developed postural orthostatic tachycardia syndrome (POTS), heart palpitations, gastroparesis, urinary incontinence, diplopia, and seizures. She also reported experiencing auditory and visual hallucinations. She underwent dysautonomia evaluation. In addition, she was recommended a full power wheelchair for the best quality of life to reduce injuries including concussions caused by frequent falls. This recent presentation has left K10031-10133 feeling overwhelmed and unsatisfied, resulting in the development of anxiety and depression diagnoses.

Her tilt table test yielded a positive result. The ophthalmic exam revealed unusual changes to her optic disks but without an elevated intraocular pressure, suggesting that her large optic nerves might represent physiologic cupping rather than glaucoma. Her brain MRI showed nonspecific findings, including a subtle focus of T2 signal abnormality involving the subcortical white matter of the right parietal lobe without associated enhancement. Other negative diagnostic test results included kidney ultrasound, chest X-ray, thyroid profile, urine vanillylmandelic acid (VMA) level, catecholamines panel (urine-free) and basic metabolic panel (BMP), and epinephrine and nor-epinephrine levels.

Her other remarkable medical history included a right hemisphere ischemic stroke at the age of 22. Causes for the stroke include the added risk of oral contraceptives (OCP) use for her irregular periods and a pre-existing patent foramen ovale (PFO). The stroke has led to residual left-side numbness, weakness and balance issues, as well as apraxia and dysarthria. Her other diagnoses include asthma, joint stiffness, hyperlipidemia, sleep walking, and dyspnea. See Table 2 for proband K10031-10133’s clinical phenotype list with HPO annotations. See supplemental note 6 for a full report of HPO analysis on proband K10031-10133. A video recording (HDV_0079) of K10031-10133 explaining her medical presentation is described in the supplemental videos section and can be provided on request to qualified investigators.

**Table 2.** Clinical Presentation of Proband K10031-10133’s Family

| **Table 2. Clinical Presentation of Proband K10031-10133's Family** |  |  |  |  |  |  |  |
| --- | --- | --- | --- | --- | --- | --- | --- |
| **Pedigree K10031** |  | **10133** | **10145** | **10235** | **10261** | **10138** | **10143** |
| **General Information** |  |  |  |  |  |  |  |
| Age (years) |  | 26 | 45 | 24 | 22 | 20 | 18 |
| Gender |  | F | F | F | M | M | F |
| **Clinical Manifestations** | **HPO#** |  |  |  |  |  |  |
| **Developmental/Growth** |  |  |  |  |  |  |  |
| Dyslexia | 0010522 | - | - | - | + | - | - |
| **Cardiovascular** |  |  |  |  |  |  |  |
| Bradycardia | 0001662 | + | - | - | - | + | - |
| PFO | 0001655 | + | - | - | - | - | - |
| Syncope | 0001279 | + | + | - | + | + | - |
| Tachycardia | 0001649 | + | + | - | - | - | - |
| **Eyes** |  |  |  |  |  |  |  |
| Astigmatism | 0000483 | + | - | - | - | - | - |
| Diplopia | 0000651 | + | - | - | - | - | - |
| Myopia | 0000545 | + | - | - | - | - | - |
| Optic disks changes | NF | + | - | - | - | - | - |
| Peripheral vision | NF | + | - | - | - | - | - |
| Prominence to the eyes | NF | + | + | + | + | + | + |
| **Gastrointestinal** |  |  |  |  |  |  |  |
| Acid reflux | 0002020 | - | - | + | - | - | - |
| Anorexia | 0002039 | - | + | - | - | - | - |
| Gastroparesis | 0002578 | + | - | - | - | - | - |
| Nausea | 0002018 | + | - | - | - | - | - |
| **Gynecologic & Genitourinary** |  |  |  |  |  |  |  |
| Irregular periods | NF | + | - | - | NA | NA | - |
| Urinary retention | 0000016 | + | - | - | - | - | - |
| Urinary incontinence | 0000020 | + | - | - | - | - | - |
| **Hematologic/Lymphatic/Immunologic** |  |  |  |  |  |  |  |
| Adenopathy | Lymphadenopathy*:<br>0002716 | + | - | - | - | - | - |
| **Musculoskeletal** |  |  |  |  |  |  |  |
| Arthralgia | 0002829 | + | - | - | - | - | - |
| Cyanotic lower extremities | 0001063 | + | - | - | - | - | - |
| Joint Stiffness | 0001387 | + | - | - | - | - | - |
| Muscle weakness | 0001324 | + | - | - | - | - | - |
| Osteoporosis | 0000939 | - | + | - | - | - | - |
| **Neurological** |  |  |  |  |  |  |  |
| Apraxia | 0002186 | + | NK | NK | NK | NK | NK |
| Arthritis | 0001369 | + | - | - | + | - | + |
| Auditory hallucinations | 0008765 | + | - | - | - | - | - |
| Concussion | NF | + | - | - | - | - | - |
| Convulsions | NF | + | - | - | - | - | - |
| Dizziness | NF | + | + | - | + | - | + |
| Dysarthria | 0001260 | + | - | - | - | - | - |
| Fatigue | 0012378 | + | + | - | - | - | - |
| Frequent falls | 0002359 | + | - | - | - | + | - |
| Headache | 0002315 | + | NK | NK | NK | NK | NK |
| Heat intolerance | 0002046 | + | + | - | - | - | - |
| Ischemic stroke | 0002140 | + | - | - | - | - | - |
| Migraine | 0002076 | + | - | + | + | + | + |
| Numbness | Paresthesia*:<br>0003401 | + | - | - | - | - | - |
| Seizure | 0001250 | + | - | - | + | - | - |
| Tremor | Postural/Resting | + | + | + | + | + | + |
|  | tremor*:<br>0002174/0002322 |  |  |  |  |  |  |
| Visual hallucinations | 0002367 | + | - | - | - | - | - |
| **Respiratory** |  |  |  |  |  |  |  |
| Asthma | 0002099 | + | + | + | + | + | + |
| Dyspnea | 0002094 | + | NK | NK | NK | NK | NK |
| **Psychiatric** |  |  |  |  |  |  |  |
| ADHD | 0007018 | - | - | - | + | + | - |
| Anxiety | 0000739 | + | + | + | - | + | + |
| Depression | 0000716 | + | + | - | - | - | - |
| Dissociative identity disorder | NF | - | + | - | - | - | - |
| Obsessive-compulsive behavior | 0000722 | - | + | - | - | + | - |
| Tourette syndrome | NF | - | - | - | - | + | - |

**Abbreviation:** NA- Not applicable, NF- Not found, NK- Not Known.
\*Terms listed in front of this symbol refer to the synonym of the disease or feature in question as categorized in the HPO Phenomizer Tool.

Her family history is remarkable for not only dysautonomia-like symptoms, but also for other conditions. Her mother and all four siblings present dysautonomia-like symptoms to some degree, such as dizziness and fainting. In addition, a younger sister (K10031-10235) has gastroesophageal reflux and anxiety. A young brother (K10031-10261) has seizures, attention deficit, dyslexia, and asthma. Another younger brother (K10031-10138) has TS. Another younger sister (K10031-10143) has asthma, anxiety, and tremor. Her mother (K10031-10145) also has HH and OCD traits. Her father has significant migraines, gastroesophageal reflux, hiatal hernia, and right sensorineural hearing loss. Other medical conditions among her first cousins include PWS, Down syndrome, cerebral palsy, TS, attention deficit hyperactivity disorder (ADHD), and bipolar disorder (Table 2). See detailed descriptions of her family members in supplemental notes. We are highlighting here that extensive characterization of families, including videotaping and the collection of collateral information from other relatives, yields a rich texture of findings that are not always easily captured in written medical records.

### Genomic analyses

We previously reported a remarkably large false negative rate with the Complete Genomics platform (O’Rawe et al. 2013a), so we chose to utilize the Illumina platform for whole genome sequencing. Nine members of the family underwent WGS, enabling a wide scope of variant calling from a single SNP to large copy number events. To reduce false variant calls, more than one pipeline was used to detect SNPs, INDELs, large SVs, and CNVs, as we previously suggested (O’Rawe et al. 2013a) (Figure 2).

**Figure 2.**
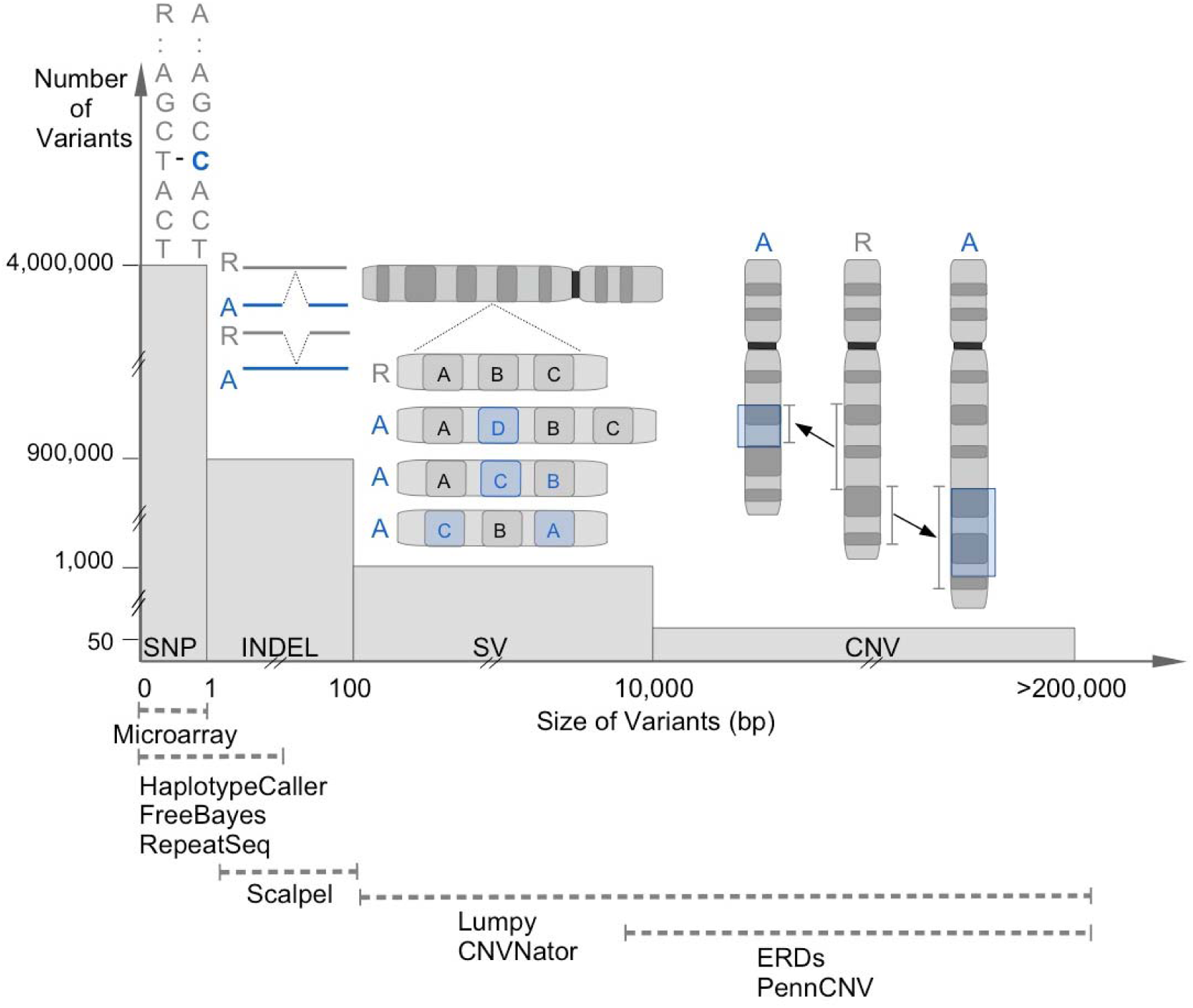
WGS can reveal a broad spectrum of variants with an integrative analysis. This is a conceptual illustration of variations in the human genome. The Y-axis shows the approximate number of variants in that category while the X-axis shows the approximate size of those variants. The interval below shows that variants of different sizes and sequence compositions can be better detected by leveraging the strength of different callers. SNP: single-nucleotide polymorphism, INDEL: insertions and deletion, SV: structural variant, CNV: copy number variants.

#### Summary statistics and quality control of the WGS data

Summary statistics for the WGS data for each sample are reported in Table 6, Table S1, Table S2, and Figure S2. The average number of reads per sample is 1,432,506,869. The number of mapped bases per sample is 124,410,724,287, with a mean coverage of the WGS data across the genome of about 40X (89% of the bases in the genome covered with at least 20X). The insert size of the libraries is about 338 and the GC content is approximately 40% across samples. With the WGS data, a mean number of 4,099,604 (SD=47,076) SNPs, 896,253 (SD=14327) INDELs, 1,284 (SD=103) SVs, and 61 (SD=4) CNVs are detected across nine samples (Table S2). Within the coding regions, the average number of SNPs, INDELs, SVs, and CNVs detected are 22,406, 2,812, 511, 12, respectively. Kinship between individuals was inferred with KING to confirm family relationship between research participants in this study (Table S3) (Manichaikul et al. 2010).

#### *Identified de novo* deletions (totaling 5.5 Mb) in region 15q11.2 to 15q13.1 in the PWS proband

ERDS and CNVnator both detected three *de novo* heterozygous deletions with a total size of about 5.5 Mb, in the chromosome regions from 15q11.2 to 15q13.1 in the male proband with PWS (10232, Figure 3). Due to the high resolution of the WGS data, the actual genomic coordinates of the breakpoints can be revealed relative to reference genome hg19; chr15:22,749,401-23,198,800 (∼449 Kb), chr15:23,608,601-28,566,000 (∼4.96 Mb), and chr15:28,897,601-28,992,600 (∼95 Kb). Notably, these deletions are very close to one another; the distances between each deletion are only ∼410 Kb and ∼332 Kb, respectively. Due to the lack of sequencing data from the proband’s mother, additional analysis was performed to determine whether the paternal allele or the maternal allele is deleted. This can be inferred through SNPs where the mendelian inheritance law is violated; meaning those instances in which the proband (K10031-10232) does not carry certain paternal or maternal SNPs that his brother (K10031-10233) does carry. In total, there are 2,987 SNPs where the proband’s father (K10031-10231) is a homozygote and the proband’s brother (K10031-10233) is a heterozygote. Out of the 2112 SNPs where the father (K10031-10231) is homozygous to the reference allele, the proband is homozygous to the alternative allele at 1944 loci (yellow-tagging SNPs in Figure 4). On the other hand, for 875 SNPs where the father (K10031-10231) does not carry any reference allele, the proband carries only the reference allele at 861 SNPs (yellow-tagging SNPs in Figure 4). This indicates that the proband only carries the maternal allele in those regions. Within the genomic regions containing the *de novo* deletions, the depth of coverage in the proband’s genome is about 20X, while the mean coverage across the genome is about 40X.

**Figure 3.**
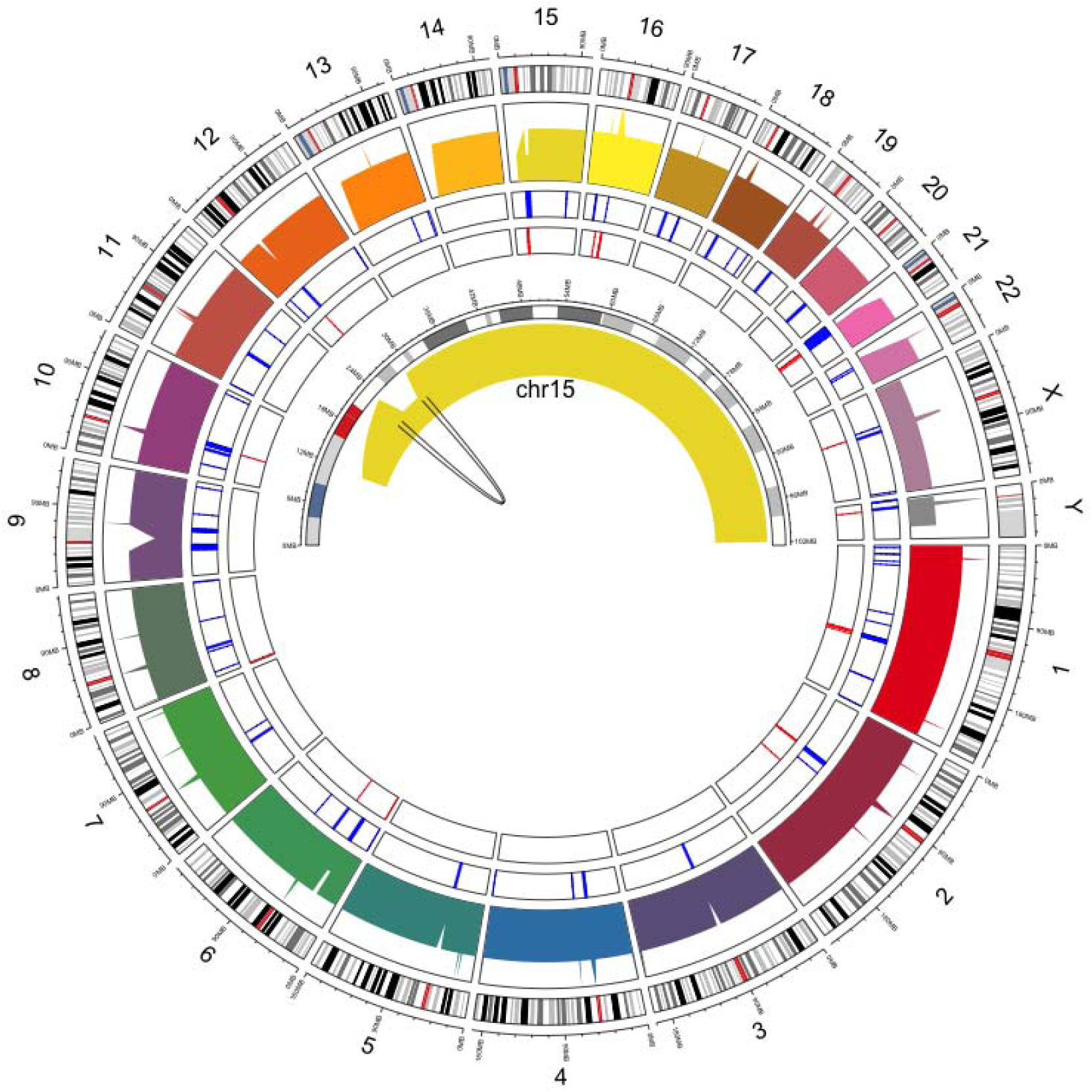
Circos plot of the PWS proband’s genome, highlighting chromosome 15. The outer circle is the cytoband of the human genome. The inner circle is the genome coverage of the PWS proband’s (K10031-10232) genome. The breakpoint of the 15q11.2-15q13 deletion region in chromosome 15 is illustrated in the center.

**Figure 4.**
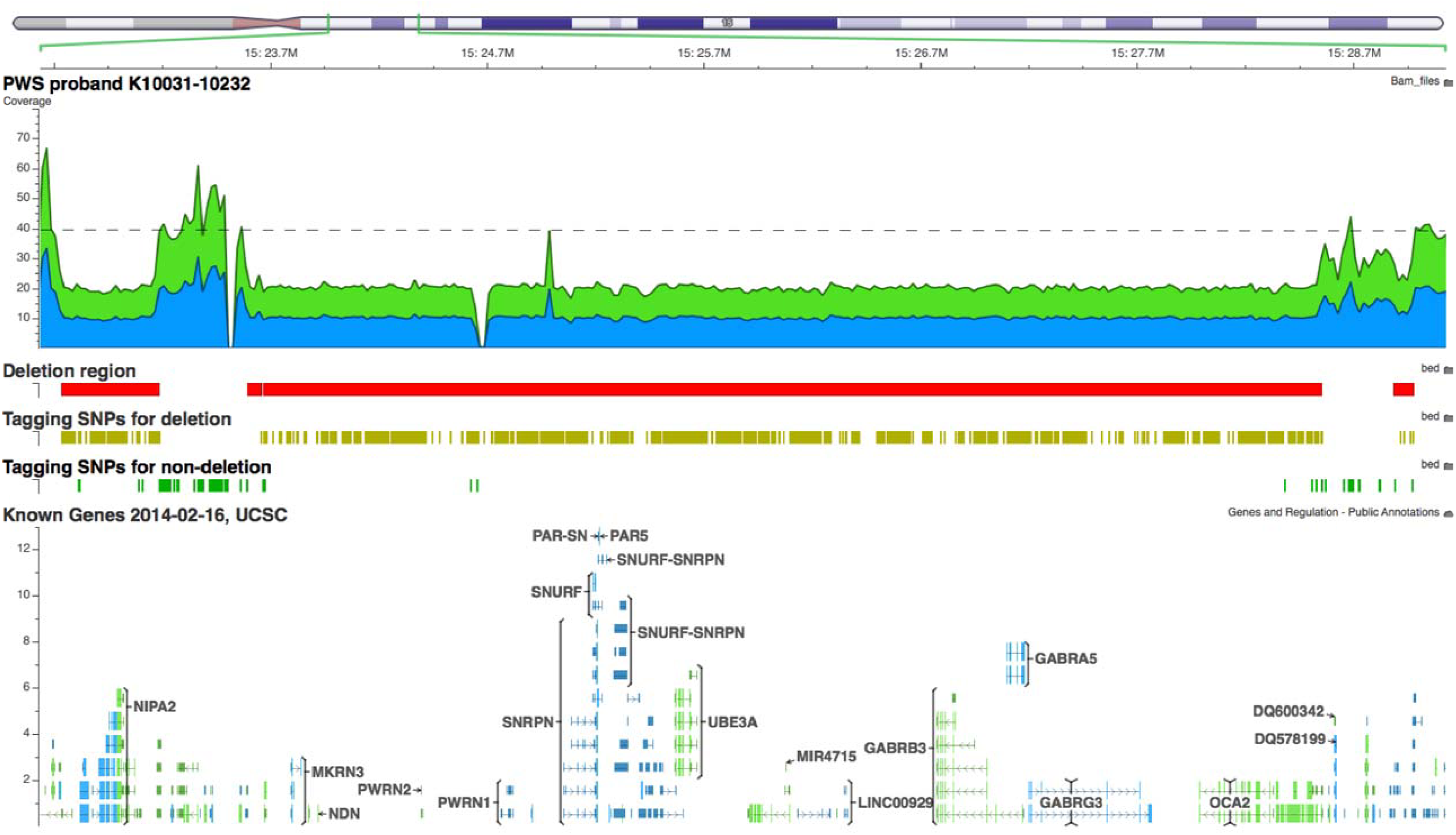
Screenshot of three heterozygous *de novo* deletions between the region 15q11.2 to 15q13 in the proband. The deleted regions in K10031-10232 are denoted by the red boxes and the tagging SNPs (yellow). The non-deleted regions are denoted by the green tagging SNPs. Genome-wide average coverage (40X) is denoted by the grey dashed line. The breakpoints of these deletions (PWS Type I deletion) are chr15:22,749,401-23,198,800 (∼449 Kb), chr15:23,608,601-28,566,000 (∼4.96 Mb), and chr15:28,897,601-28,992,600 (95 Kb) (hg19). These deletions are not detected either in the proband’s father or the unaffected brother, and were confirmed with the Illumina Omni 2.5m microarray data.

These deletions were not detected in either the proband’s father or brother using the WGS data (Figure S7). The orthogonal Illumina microarray data further confirmed this discovery; his father (K10031-10231) and his brothers (K10031-10233 and K10031-10234) do not carry any of these deletions in their genome (Figure S3, S5, S6). Probe distributions of Log-R ratios (LRR) and B allele frequencies (BAF) are not uniform in the microarray because the density of SNP varies between genomic regions (Figure S3 – Figure S6). This highlights the higher resolution and completeness of WGS over microarray for precise molecular diagnosis of such diseases. Using other methods would make it necessary to go through a more complicated workflow with standard genetic testings for PWS (Dittrich et al. 1992; Cassidy and Driscoll 2008; Cassidy et al. 2012). Since Angleman Syndrome (AS) and PWS share a similar cytogenetic anomaly in 15q11.2 to 15q13 (Knoll et al. 1989; Nicholls and Knepper 2001), WGS could potentially help reveal the sub-types of both syndromes because the breakpoints of CNVs can be mapped at nucleotide level and one could distinguish which allele (paternal or maternal) has been deleted if the WGS data from the parents is also provided. However, WGS would not be enough to detect either uniparental paternal disomy or imprinting defect in this genomic region for non-deleted PWS individuals (Malcolm et al. 1991; Cassidy et al. 2012).

This study confirms that the proband carries the PWS Type I deletion (spanning breakpoint BP1 and distal breakpoint BP3) defined by previous publications (Christian et al. 1995; Cassidy et al. 2012). Additionally, the second deletion in this proband (K10031-10232) embraced a highly restricted deletion (∼118 Kb) reported previously in an individual with PWS (Bieth et al. 2014). The complete list of genes that fall into the deletion regions are described in Table 3. On the other hand, the coordinates for the non-deleted region is chr15: 23383801-23679100. The genes that fall in this non-deleted region are mostly pseudogenes, small RNAs, and long non-coding RNA (lncRNA) genes, including *WHAMMP3* (partially deleted, exon 1 to exon 5), *GOLGA8IP, DQ600342, DQ582939, DQ578838, DQ588973, DQ595055, DQ572979, JB175342, HERC2P2, DQ582073, LOC440243, DQ594309, DQ595648, GLOGA8EP, DQ600342, DQ578838, DQ572979, JB175342, LOC440243, DQ582073, DQ595648, GOLGA8S,* and *DQ578199*.

**Table 3.** A list of genes that fall into the deletion regions. The break points are chr15:22,749,401-23,198,800, chr15:23,608,601-28,566,000, and chr15:28,897,601-28,992,600. The coordinates are shown with respect to hg19. All genes are heterozygous for the deletion.

| Gene category | Genes |
| --- | --- |
| Protein coding genes | *TUBGCP5, CYFIP1, NIPA2, NIPA1, MKRN3, MAGEL2, NDN, NPAP1, SNRPN, SNURF, SNURF-SNRPN, HBT8, C15ORF49, UBE3A, ATP10A, GABRB3, GABRB5, GABRG3, OCA2, HERC2* |
| Pseudogenes | *GOLGA8S, GLOGA8EP, GOLGA6L2, GOLGA8IP, WHAMMP3* (partially deleted, exon 1 to exon 2), *HERC2P2, AK124131, AX747189, AK124673, HERC2P9, WHAMMP2* (partially deleted, exon 1 to exon 5) |
| microRNAs | *MIR4508, MIR4715* |
| piRNAs | *DQ600342, DQ582939, DQ578838, DQ588973, DQ595055, DQ572979, DQ582073, DQ594309, DQ595648, DQ578199, DQ596685, DQ582448, DQ593032, DQ588687, DQ597560* |
| snoRNAs | *SNORD109B, SNORD116- (1, 2, 3, 4, 5, 8, 10, 11, 12, 13, 15, 16, 18, 19, 20, 22, 24, 25, 26, 28, 29), SNORD115- (1, 2, 3, 4, 5, 6, 7, 8, 9, 10, 11, 13, 14, 15, 16, 17, 22, 24, 25, 26, 27, 28, 30, 31, 32, 33, 35, 37, 38, 39, 40, 41, 44, 45, 47, 48)* |
| Non-coding RNAs (ncRNA) | *PWRN2, PWRN1, PWARSN, PWAR1, PWAR5, LINC00929, LOC100128714, LOC283685, LOC440243, LOC283683* |
| tRNA | *tRNA_Glu* |
| Unassigned RNA | *JB175342* |

#### Identified p.C282Y variant in the mother with HH and other unrelated findings

The mother (K10031-10145) with HH is homozygous for the p.C282Y variant in HFE, which is consistent with her molecular genetic assay results. Results from analyzing the WGS data showed that her brother (K10031-10231) is also homozygous for the p.C282Y variant in HFE (Table 4). However, his clinical test result has not provided any evidence to support the diagnosis of HH so far, even though male p.C282Y homozygotes are more likely to develop iron-overload–related diseases due to the lack of the iron clearance events such as menstruation and pregnancy in women (Allen et al. 2008). This is in line with the well-known fact that even family members can have variable expressivity of disease, including not having any detectable symptoms. There are many publications showing that the phenotypic expression of a given mutation in *HFE* may vary widely (Hanson et al. 2001; Pietrangelo 2004). Some studies previously estimated that less than 1% of individuals in the U.S. carrying homozygous mutations present clearly with clinical diagnosis of hemochromatosis (Beutler et al. 2002). In contrast to studies that have searched for the “causal” gene, some have reported that genetic variations can instead have large effects on phenotypic variability, suggesting underlying genetic complexity from multiple interacting loci (Mackay et al. 2009; Massouras et al. 2012; Zuk et al. 2012; Corbett-Detig et al. 2013). Understanding such diseases thus requires probabilistic thinking about the risk of developing the clinical manifestation, rather than deterministic genotype-phenotype “causation” (Moczulski et al. 2001; Thornton-Wells et al. 2004; Freund et al. 2013; Lyon and O’Rawe 2014).

**Table 4.** A list of variants with previous evidence in ClinVar found in the pedigree members. The coordinates are shown with respect to hg19. The mother with hereditary hemochromatosis is homozygous for the p.C282Y variant in HFE. The carriers are represented by the numbers shown in Figure 1.

| Gene | Genomic coordinates | Change & variant type | Zygosity & Carriers | AAF | Relevant diseases & Inheritance |
| --- | --- | --- | --- | --- | --- |
| **HFE** | chr6:<br>26093141 | G>A<br>missense | hom: 10145, 10231<br>het: 10232, 10233,<br>10133, 10235,<br>10138, 10143 | 0.007% | hereditary<br>hemochromatosis -<br>AR) |
| **BRIP1** | chr17:<br>59937223 | G>C<br>missense | het: 10231, 10145,<br>10133, 10235,<br>10138 | 0.04% | Breast cancer,<br>early-onset - AR |
| **MKKS** | chr20:<br>10393439 | G>T<br>missense | het: 10133, 10145,<br>10143 | 0.9% | Mckusick-Kaufman<br>Syndrome - AR |
| **PRSS1** | chr7:<br>142458451 | A>T<br>missense | CH: 10231, 10232,<br>10145, 10143,<br>10133, 10235,<br>10138 | 47% | hereditary<br>pancreatitis - CH |
|  | chr7:<br>142458526 | A>G<br>missense |  | 3% |  |

\*AAF: Alternate Allele Frequency in ExAC database, Hom: homozygous, het: heterozygous, AR: Autosomal recessive, CH: Compound heterozygous.

As previously argued, there is nothing ‘incidental’ about unrelated findings (Lyon 2012b). Regardless of how one defines such terms, it is highly likely to find genetic evidence that might be unrelated to the primary research purpose (Christenhusz et al. 2013). And, this genetic information can inform participants’ clinical treatments, which may not have been taken into consideration previously (O’Rawe et al. 2013d). Thus, alongside the primary research-focused analysis, the participating subject and family also received the research findings (Table 4). Seven individuals (K10031-10231, K10031-10232, K10031-10145, K10031-10133, K10031-10235, K10031-10138, K10031-10143) in this pedigree are compound heterozygous for variants c.86A>T and c.161A>G in *PRSS1*, which were previously reported to be associated with hereditary pancreatitis (Teich et al. 2005). In addition, five people (K10031-10231, K10031-10145, K10031-10133, K10031-10235, K10031-10138) were found to be carriers for c.139C>G variant in *BRIP1*, which was previously associated with early-onset breast cancer (Cantor et al. 2001). Three people (10133, 10145, 10143) are carriers for the c.724G>T variant in *MKKS*, which was reported to be associated with a rare autosomal recessive developmental anomaly syndrome, namely McKusick-Kaufman syndrome (Stone et al. 2000).

#### In search of variants that might contribute to dysautonomia-like symptoms

The findings above identified two known variants that were previously reported in the literature, suggesting they are likely large effect-size variants contributing to PWS and HH. On the other hand, none of the family members with dysautonomia-like symptoms carry any previously reported variants in *IKBKAP* that are implicated in the autosomal recessive transmission of FD, which is also called Riley–Day syndrome and hereditary sensory and autonomic neuropathy type III (HSAN-III). The WGS data have effective sequence coverage (> average coverage 40X) for this gene but no novel rare variants in either the coding regions or the intronic regions were identified. It is worth noting that both the mother (K10031-10138) and one of her brothers (K10031-10145) carry heterozygous variants of p.H604Y and p.G613V in the protein product of *NTRK1*, which has been proven to contribute to HSAN IV (congenital insensitivity to pain with anhidrosis (CIPA). CIPA is a disease closely resembling FD (HSAN III), and is characterized by a lack of pain sensation, anhidrosis, unexplained fever since childhood, self-mutilating behavior, and intellectual disability of varying degree (Swanson 1963; Indo et al. 1996). However, since both variants have been reported before in CIPA individuals diagnosed with cancer as well as healthy individuals, they are considered to be polymorphisms in the population (Cargill et al. 1999; Gimm et al. 1999; Shatzky et al. 2000; Indo 2001; Greenman et al. 2007). These two variants seem to be linked since they always seem to occurr together (Gimm et al. 1999). Both variants are located within the intracellular tyrosine kinase domain (amino acids 510-781) of the encoded protein. However, both sites are not conserved and biochemistry studies further confirmed that neither of these two variants has any effect on protein expression and phosphorylation compared to the wild type (Mardy et al. 2001). The fact that the mother’s brother (K10031-10231) also carries these two variants further proved that they are likely to be polymorphisms, and neither of these variants is present in the proband (K10031-10133). Hence, we investigated whether the dysautonomia-like symptoms presenting within the family are possibly arising in conjunction with other mutations. To investigate, we used pVAAST, CADD, and other prioritizing tools to help leverage the power of the large pedigree and WGS. Table S4 summarizes the lists of variants that meet the following criteria: 1) found in the probands and not found in unaffected people in the family, 2) called by at least one pipeline and supported by a second pipeline, 3) located within coding regions, 4) have high rankings by pVAAST and with either a Combined Annotation Dependent Depletion (CADD) score greater than 15 or at least with medium effect predicted by GEMINI, and 5) have Alternate Allele Frequency (AAF) < 1% in either ExAC and 1000G databases.

#### Pharmacogenomic analyses for individual K10031-10133

Pharmacogenomic analyses were performed on individual K10031-10133 using the Omicia Opal platform, which is based on dosage-calculating algorithms from the PharmGKB database (Hewett et al. 2002). Pharmacogenetic variants influencing the metabolism of warfarin and simvastatin were found. These two medications were being routinely prescribed to K10031-10133. Pharmacogenomic analysis guidelines and algorithms were obtained from the International Warfarin Pharmarcogenomics Consortium (IWPC) and the Clinical Pharmacogenomics Implementation Consortium (CPIC) (The International Warfarin Pharmacogenetics 2009; Johnson et al. 2011; Wilke et al. 2012). Dose calculation of Walfarin was done based on the IWPC algorithm, which considers the individual’s *CKORC1* and *CYP2C9* genotype, age, height, weight, race, history of enzyme inducer and amiodarone (Supplemental data 4). The calculation suggests a warfarin dosage at 5.85 mg per day warfarin for 10031 (Table 5). For simvastatin, the CPIC algorithm suggests modestly-increased myopathy risk for people with the C allele at *SLCO1B1* rs4149056 taking simvastatin doses of 40 mg daily (Table 5). Thus, a lower dosage of simvastatin at 20 mg per day is recommended instead.

**Table 5.**
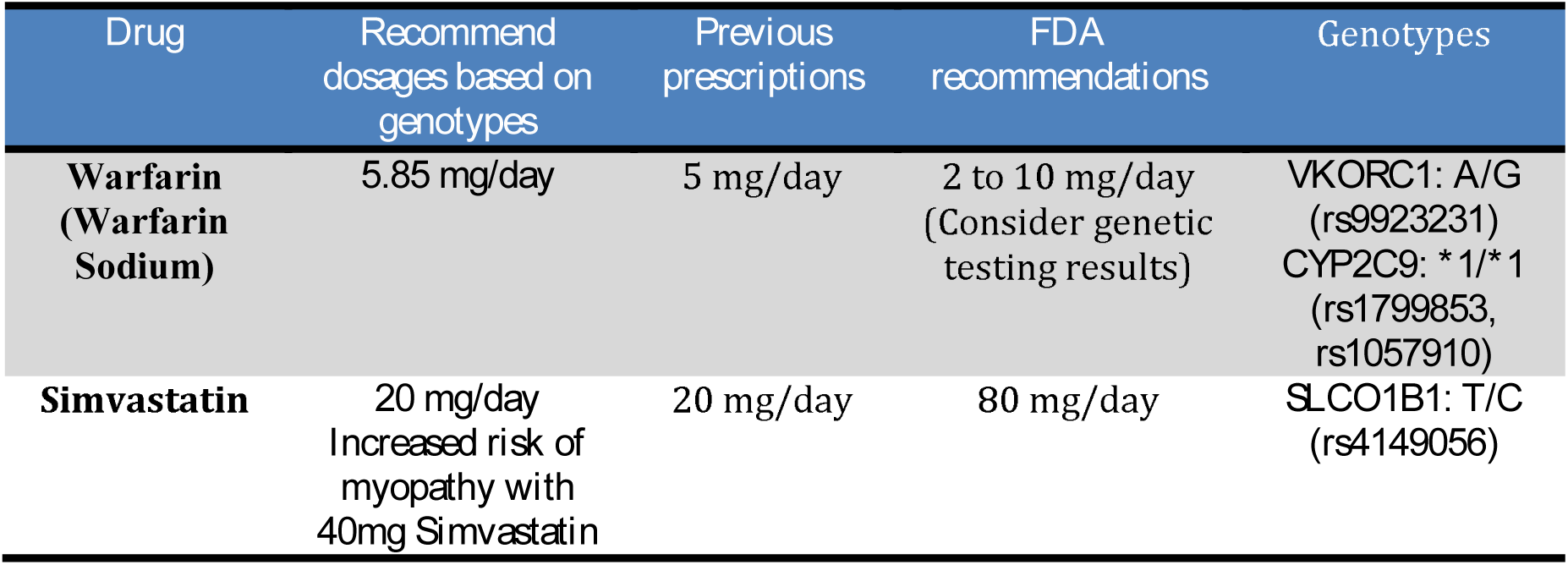
Recommended dosages for warfarin and simvastatin dosages based on the individual K10031-10133’s SNP results from the WGS data in comparison to what she was actually prescribed in the absence of any genetic testing. Pharmacogenomics analyses were performed based on guidelines and algorithms from the International Warfarin Pharmarcogenomics Consortium (IWPC) and the Clinical Pharmacogenomics Implementation Consortium (CPIC) in the PharmGKB database. People who are homozygous for major alleles at both sites in *CYP2C9* are designated as *1/*1.

| Drug | Recommend dosages based on genotypes | Previous prescriptions | FDA recommendations | Genotypes |
| --- | --- | --- | --- | --- |
| **Warfarin (Warfarin Sodium)** | 5.85 mg/day | 5 mg/day | 2 to 10 mg/day<br>(Consider genetic testing results) | *VKORC1*: A/G (rs9923231)<br>*CYP2C9*: *1/*1 (rs1799853, rs1057910) |
| **Simvastatin** | 20 mg/day<br>Increased risk of myopathy with 40mg Simvastatin | 20 mg/day | 80 mg/day | *SLCO1B1*: T/C (rs4149056) |

The resulting calculation is very close to the previous prescriptions from the individual’s cardiologist, although the pharmarcogenomic recommendation for warfarin is slightly higher than the actual prescription (0.85 mg/day). This suggests that for this single individual, pharmacogenomics results confirm the appropriate dosages for these two medications. In fact, the FDA recommended doses for warfarin consist of a wide range, namely 2 to 10 mg per day. For simvastatin, the recommended FDA dose is 80 mg daily, which is much higher than the actual prescription dosages. In this case, dosage calcuation purely relied on the individual’s genotype and other general information. One could include this information at least for a genotype-driven prescription.

## DISCUSSION

In conjunction with a previous report by Schaaf et. al., this research report provides insight into using WGS as a genetic test to investigate a well-known syndrome, Prader-Willi syndrome (Schaaf et al. 2013). Notably, this is the first report of an Illumina HiSeq WGS experiment on an individual with PWS with the paternal deletion of 15q11.2-q13. One group previously used the Complete Genomics platform to detect *de novo* small frameshift deletions in *MAGEL2*, a gene within the known PWS genomic domain, although 65–75% of PWS is due to the paternal deletion of 15q11.2-q13 (Cassidy et al. 2012; Schaaf et al. 2013). In our study, three *de novo* deletions were discovered at single base pair resolution and provide additional evidence for more complex recombination events in the chromosome region 15q11.2 to 15q13. Compared to WGS, CNV detection at a genome-wide scale on Illumina 2.5M microarray were not as sensitive due to uneven distribution of SNP target probes. DNA methylation analysis will detect more than 99% of PWS individuals, but additional genetic tests are necessary to define the molecular sub-types (Cassidy et al. 2012). Thus, WGS along with methylation analysis should provide a comprehensive view of PWS diagnosis support. However, identifying disease-relevant loci is just the beginning of studying any human diseases. Downstream functional follow-up experiments are essential to unlocking and understanding the underlying mechanism and for subsequent development of therapeutic strategies. For example, thorough characterization of patient-derived induced pluripotent stem cells (iPSCs) successfully revealed the cross-talk between the PWS and DLK1-DIO3 loci in early human development and the under-appreciated complex phenomenon in parental imprinting (Stelzer et al. 2014). Focused studies could also faciliate the understanding of previously unannotated class of lncRNAs in the molecular pathogenesis of PWS (Yin et al. 2012).

The HPO Phenotype Ontology Phenomizer tool is innovative in that it provides a standardized vocabulary bank of phenotypic abnormalities to describe components and presentations of human pathologies. Others have previously reported that phenotypic matching can help interpret CNV findings based on integrated cross-species phenotypic information (Köhler et al. 2014). The paper’s usage of it in the form of a clinical diagnostics implement proves advantageous in that it sources from a wealth of medical literature and databases such as Online Mendelian Inheritance in Man (OMIM) (Robinson et al. 2008). During the selection process for a particular patient’s feature, one is thus able to acquire a surplus of clinical and scientific knowledge about the diseases linked to the feature in question, which includes descriptions, phenotype-gene relationships, clinical features, inheritance, molecular genetics, etc. This aptly aids one in selecting for the most appropriate description that fits the patient. Following this collection of features is the ability to acquire a clinical diagnosis, which is helpful in considering possible diagnosis and treatment (see supplemental notes for examples of full HPO Phenomizer Diagnosis Reports). As the field of medical genetics advances, researchers will need to find an efficient way to capture phenotypic information that allows for the use of computational algorithms to search for phenotypic similarity between genomics studies (Robinson and Mundlos 2010).

For the study on the family members with dysautonomia-like symptoms, the unambiguous description of their physical symptoms and the incompleteness of their clinical investigation and records hinder both the analysis of their phenotypic data and the interpreation of their genotypic data. As a matter of fact, an inaccurate phenotypic data can be quite misleading in finding the disease-related variants, since often times construction of a disease inheritance model to partition the variants is necessary, and is largely based on the segragation of the phenotype in the pedigree of study. In this particular family, the inheritance model seems to be a dominant pattern since multiple family members in different generations share similar symptoms such as dizziness and syncope. However, such assumptions might not necessarily to be true due to the unclear segregation of more specific and objective clinic findings like absence of gungiform papillae of the tongue and decreased deep tendon reflexes for FD and anhidrosis for CIPA. To further facilitate and pin down the disease-relevant variants in this family, emphasis needs to be placed on performing more clinical investigations including diagnostic tests and collecting the complete medical records from as many family members as possible.

The consequence of general and incomplete medical notes is not limited to discovering the variants; it also inhibits some features from being selected for and included in the HPO analysis. For instance, K10031-10133 is reported to have “irregular menses”, but this term does not appear when put into the “features” search tab or “disease” search tab directly. Instead, the feature was pursued under the “ontology” tab, which is a top-down approach to selecting features. The available features relating to an irregular menses in the tool include amenorrhea, delayed menarche, menometrorrhagia, menorrhagia, etc., but the proband’s medical documents lacked the details necessary for selecting the best option. However, this kind of information is necessary for more accurate, optimal Phenomizer results, and emphasizes the necessity of data availability as well as clear descriptions of the affected persons in question for phenotype analysis to achieve its full potential (Robinson and Mundlos 2010).

The Phenomizer diagnostic results for the probands, K10031-10133 and K10031-10232, highlight the need for a more flexible method of inputing features as well as adjustments for each feature in influencing the diagnostic results. Previous literature suggests that usage of complete features has lead to more accurate diagnosis results (Zemojtel et al. 2014). This was the case for K10031-10232, for whom there was a plethora of clinical data available, so that selecting 10232’s features proved to be a smoother process with more precise results. A combination of more standardized language in the clinical setting, as well as detailed, accurate reports of the patient’s medical history, is necessary to select for accurate features in the HPO tool. The Phenomizer tool’s vocabulary bank may also seek to build a wider range of descriptions, and the tool’s searching algorithm may be developed to better incorporate the vague inquiries described above. Potential remains for development of a more multi-dimensional depiction of subjects that takes into account the past and present human presentation, and will aid in efforts for early diagnoses and intervention.

## METHODS

### Enrollment of research participants

The collection and analysis of DNA was conducted by the Utah Foundation for Biomedical Research, as approved by the Institutional Review Board (IRB) (Plantation, Florida).

### Clinical phenotyping of individuals participating in this study

The family was interviewed by one of us (GJL), who is a board-certified child, adolescent and adult psychiatrist. Medical records were obtained and reviewed, in conjunction with further interviews with the family. The interviews were videotaped and later reviewed to facilitate further diagnostic efforts.

### Clinical evaluation/diagnostics

Various clinical diagnostic testing was performed on proband K10031-10133, including tilt table test, brain MRI, ultrasound of the kidneys and chest X-ray. In addition, her cholesterol level, thyroid profile, urine vanillylmandelic acid (VMA), catecholamines panel (urine-free), basic metabolic panel (BMP), and epinephrine and norepinephrine levels were also screened. For proband K10031-10232, the following diagnostic testing was performed: multiple sleep latency test (MSLT), autism diagnostic observation system (ADOS) - module 2, the Childhood Autism Rating Scale (CARS), Behavior Assessment System for Children (BASC), Intelligence Quotient (IQ), Abnormal Involuntary Movement Scale (AIMS), as well as electrocardiogram (EKG), polysomnographic report, and echocardiogram.

### HPO analysis of the phenotypes

HPO analyses were conducted for both probands 10232 and 10133. Their clinical features were inserted into the HPO Phenotype Ontology Phenomizer clinical diagnostics tool using the “Any” mode of inheritance for as complete coverage of possible diagnoses and inheritance patterns as possible. The Resnik (not symmetric) mode of analysis was also selected. The features are listed and integrated in Table 1 and 2, and complete Phenomizer Diagnosis forms are available in the supplemental portion of the paper. There are different methods of searching for the proper HPO Phenomizer feature. One can search for the feature directly by inserting it into the “Features” tab, look for it as associated with a disease via the “Diseases” tab, or locate it via the “Ontology” tab by selecting for the organ abnormalities that result in that particular feature. This last one is a top-down approach in that one selects the proper organ involved, then the system that is affected, followed by abnormality in morphology or physiology, etc., making one’s search more and more specific until one locates the desired feature. Following this collection of features is the ability to acquire a clinical diagnosis, which is helpful in considering possible diagnosis and treatment.

### Generation of WGS and microarray data

Blood and saliva samples were collected from nine individuals from the extended pedigree described in the results. Two CLIA-certified WGS tests (K10031-10133 and K10031-10138) were performed at Illumina, San Diego. The other seven WGS were performed at the sequencing center at Cold Spring Harbor Laboratory (CSHL). All of the libraries were constructed with PCR amplification. All libraries were sequenced on Illumina HiSeq2000 with average paired-end read length of 100 bp. Since the DNA extracted from saliva samples contains a certain proportion of bacterial DNA, these samples were sequenced on additional lanes to achieve an average coverage of 40X after removing unmapped reads (Table S1). Microarray data for the same samples was generated with the Illumina Omni 2.5 microarray at the Center for Applied Genomics Core of the Children’s Hospital of Philidephia (CHOP). Illumina Genome Studio was used to extract the SNP calls and log-R ratio (LRR) from the microarray data. Finally, Qualimap (v2.0) was used to perform QC analysis on the alignment files (García-Alcalde et al. 2012). The general analysis work-flow is shown in Figure S1.

### Alignment of the short reads from WGS data

All of the unmapped raw reads were excluded to remove the sequence reads coming from the bacterial DNA (step 2 of Figure S1). The remaining reads were aligned to human reference genome (build hg19) with BWA-mem (v0.7-6a) (Li 2013). In parallel, five genomes were aligned with NovoAlign (v3.00.04) to reduce false negatives resulting from alignment artifacts. All of the alignments were sorted with SAMtools (v0.1.18) and PCR duplicates marked with Picard (v1.91) (Li et al. 2009). For the BWA-MEM bam files, INDELs were realigned with the GATK IndelRealigner (v2.6-4) and base quality scores were recalibrated (McKenna et al. 2010). For variant calling with FreeBayes, the alignment files were not processed with INDEL-realignment and base quality recalibration as these additional steps are not required by FreeBayes.

**Table 6.** Sequencing coverage for each sample in this study. A full description of the summary can be found in Table S1.

| Sample in K10031 | Number of reads | Mean coverage | Fraction covered >20X | GC content |
| --- | --- | --- | --- | --- |
| 10133 | 1,762,018,294 | 54.78X | 90.16% | 39.31% |
| 10138 | 1,413,986,946 | 44.14X | 89.31% | 39.73% |
| 10143 | 1,627,071,352 | 39.93X | 89.66% | 40.09% |
| 10144 | 1,964,930,229 | 42.63X | 89.05% | 40.08% |
| 10145 | 1,213,529,960 | 37.9X | 89.3% | 39.92% |
| 10231 | 1,204,598,812 | 37.51X | 87.54% | 40.06% |
| 10232 | 1,206,286,495 | 37.67X | 87.58% | 40.14% |
| 10233 | 1,215,246,598 | 37.9X | 87.83% | 40.34% |
| 10235 | 1,614,404,561 | 39.59X | 89.7% | 40.62% |

### Detection of SNP/INDEL/SV/CNV from the WGS and microarray data

To reduce false positive and negative variant calls, more than one pipeline was used to detect SNPs, INDELs, SVs, and CNVs. The workflow of this section is described from step 3 to step 5 in Figure S1. First, SNPs and INDELs were jointly called from nine genomes with GATK HapolotypeCaller (v3.1-1) from the BWA-MEM alignment following best practices (DePristo et al. 2011). Second, a default parameter setting was used to call variants using FreeBayes from the NovoAlign alignment (Garrison and Marth 2012). Third, Scalpel (v0.1.1) was used with the BWA-MEM bam files to identify INDELs in the exonic regions with sizes up to 100 bp (Narzisi et al. 2014). Each exon was expanded by 20 bp upstream and 20 bp downstream to reveal possible INDELs harboring splicing sites. Following the benchmarking results as recently reported (Fang et al. 2014), Scalpel INDEL calls were filtered out if they have a alternative allele coverage less than five and a Chi-Square socre greater than 10.8. Fourth, RepeatSeq (v0.8.2) was utilized to detect variants near short tandem repeat regions in the genome using default settings (Highnam et al. 2013). Fifth, Lumpy (v 0.2.6) and CNVnator were both used to call SVs with sizes ranging from 100bp and up (Abyzov et al. 2011; Layer et al. 2014). Among Lumpy calls, events supported by more than 50 reads or less than 4 reads were excluded because regions of either too low or high coverage are more likely to contain biases in sequencing or alignment. Sixth, ERDS (v1.1) was used to call CNVs from the BWA-mem bam files with default settings (Zhu et al. 2012). Among ERDS calls with a confidence score >300, duplications with sizes less than 200 Kb and deletion calls with sizes less than 10 Kb were excluded from down stream analysis. CNVnator (v0.3) was used to identify smaller CNVs that are present in the WGS data using the parameters –his 100, -stat 100, -partition 100, -call 100 (Abyzov et al. 2011). Sixth, to achieve high confidence CNV calls, PennCNV (2011Jun16 version) was used to call CNVs from the microarray data (Wang et al. 2007). Each CNV was supported by at least 10 markers, excluding CNVs with an inter-marker distance of >50 Kb. SVs and CNVs that overlapped with segmental duplication regions by 50% were also filtered out with Bedtools (Quinlan and Hall 2010).

### Filtering and annotations of the variants

To annotate the variants of interest, GEMINI (v0.11.0), ANNOVAR (2013Aug23 version), and some customized python scripts were used (step 6 of Figure S1) (Wang et al. 2010; Paila et al. 2013). The circos plot of K10031-10232’s genome was generated using circlize in R (Gu et al. 2014). The population allele frequencies (AF) were loaded with GEMINI from the 1000G database (http://www.1000genomes.org/) and Exome Aggregation Consortium (ExAC) database (http://exac.broadinstitute.org/). GEMINI also served to import the CADD C-scores, loss-of-function variants defined by LOFTEE, and the reported pathogenicity information from the ClinVar database (MacArthur et al. 2012; Kircher et al. 2014). In addition, pVAAST helped prioritize and rank the variants that might be related to certain phenotypes in this family (Hu et al. 2014).

There were several steps in filtering variants with respect to segregating patterns, population frequency, allele deleteriousness prediction, and ClinVar annotations. First, variants were partitioned by the following disease inheritance models: autosomal dominant, autosomal recessive, *de novo*, compound-heterzygous, and X-linked dominant. Second, autosomal or X-linked dominant and *de novo* variants were excluded if they had an AAF greater than 0.01 in either ExAC or 1000G database while the cut-off was increased to 0.05 for autosomal recessive and compound-heterzygous variants. Third, only the variants that met the following criteria were considered in the downstream analysis: 1) called by at least one pipeline and validated with a second pipeline, 2) ranked high by pVAAST, 3) defined as medium or high impact by GEMINI, or defined as loss of function by LOFTEE, 4) with a CADD c-score greater than 15. Fourth, variants that were considered as pathogenic, probably-pathogenic, mixed, or drug-response in the ClinVar database were also searched for. Lastly, the VCF files were also uploaded to the Omicia Opal platform and the Tute Genomics platform for online annotation, filtering, and pharmacogenomic analysis. The Tute Genomics variant interpretation report for each individual can be found in Supplemental data 5.

## List of abbreviations used

(SNP): Single-nucleotide polymorphism
(CNV): copy number variantion
(INDELs): insertions and deletions
(SV): structual variant
(WGS): whole genome sequencing
(WES): whole exome sequencing
(NGS): next-generation sequencing
(base pair): bp
(kilo base pairs): Kb
(megabase pair): Mb
(polymerase chain reaction): PCR
(PWS): Prader–Willi syndrome
(HH): hereditary hemochromatosis
(FD): familial dysautonomia
(CIPA): congenital insensitivity to pain with anhidrosis
(TS): Tourette syndrome
(HPO): the Human Phenotype Ontology
(OCD): obsessive-compulsive disorder

## ADDITIONAL INFORMATION

### Data Deposition and Access

All of the sequence reads can be downloaded under project accession number [XXXX] from the Sequence Read Archive (http://www.ncbi.nlm.nih.gov/sra).

Please note that the data have been submitted and we are awaiting deposition at SRA.

### Online Resources

The Human Phenotype Ontology (HPO): http://www.human-phenotype-ontology.org/

1000G database: http://www.1000genomes.org/

Exome Aggregation Consortium (ExAC): http://exac.broadinstitute.org/

ClinVar database: http://www.ncbi.nlm.nih.gov/clinvar/

### Ethics Statement

The collection and analysis of the DNA used in this study was conducted by the Utah Foundation for Biomedical Research, as approved by the Institutional Review Board (IRB) (Plantation, Florida). All participants in the study provided informed written consent and research was carried out in compliance with the Helsinki Declaration.

## Authors’ contributions

H.F. analyzed the sequence data. H.F., Y.W., M.Y. and G.J.L. wrote the manuscript. Y.W. and M.Y. analyzed the clinical data. Y.W. performed the sanger sequencing validation experiment and assisted in the sample preparation for the WGS. M.Y. performed the HPO analysis. J.R., L.J.B., G.H., and D.M. helped analyze the WGS and microarray data. G.J.L. supervised the data analysis. All of the authors have read and approved the final manuscript.

## Funding

The laboratory of G.J.L. is supported by funds from the Stanley Institute for Cognitive Genomics at Cold Spring Harbor Laboratory (CSHL). The CSHL genome center is supported in part by a Cancer Center Support Grant (CA045508) from the NCI.

## Competing Interests

G.J.L serves on advisory board for GenePeeks, Inc. and Omicia, Inc. D.M. was until recently an employee and shareholder of Gene By Gene, Inc. and is now Chief Scientific Officer of Tute Genomics, Inc.

## FIGURE LEGENDS

Figure legends are now beneath each figure and will be moved here later.

### Supplemental data 1

**Figure S1.**
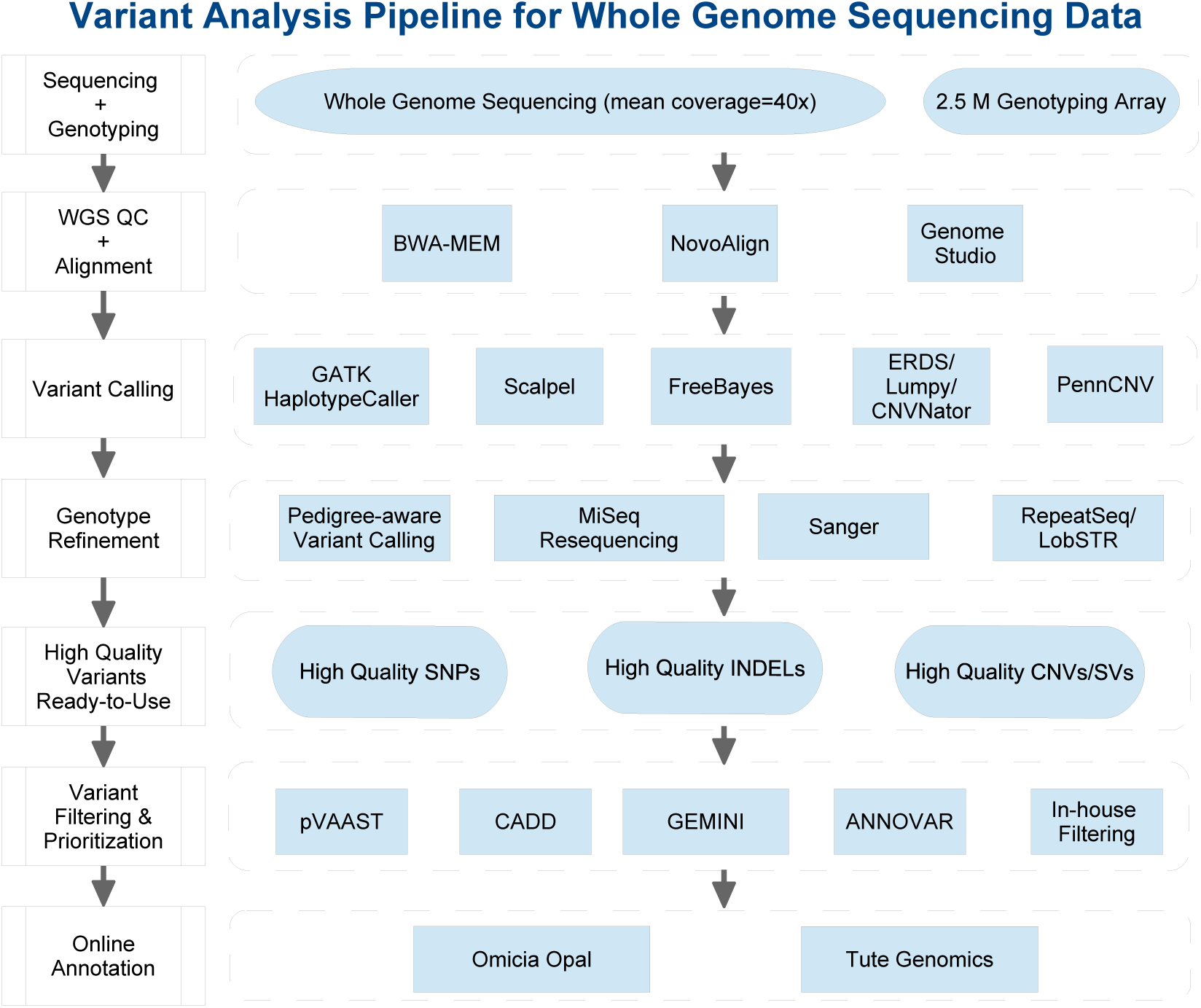
Variant analysis pipeline for whole genome sequencing data and the microarray data. The left-hand side shows the major analysis work-flow while the right-hand side shows the details of each procedure.

**Figure S2.**
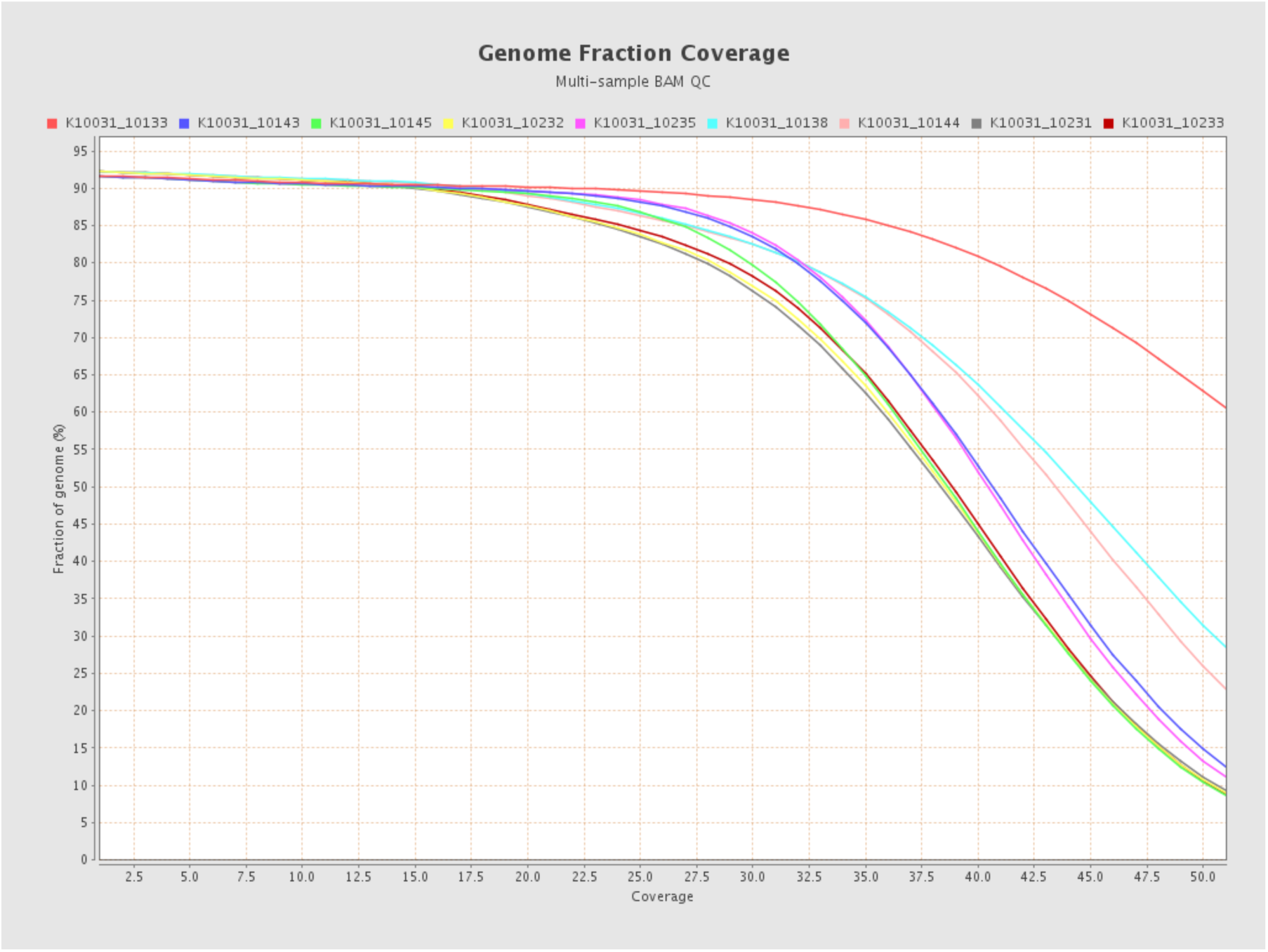
Genome fraction coverage distributions of the sequencing data. Each curve represents one genome in this study. For each sample, more than 90% of the genome is covered with about 20 reads.

**Table S1.** Summary statistics of the whole genome sequencing data generated in this study.

| Sample in K10031 | Number of reads | Percentage mapped, paired | # of mapped bases | Mean coverage | Fraction covered >20X | Median insert size | Mean mapping quality | GC content |
| --- | --- | --- | --- | --- | --- | --- | --- | --- |
| 10133 | 1,762,018,294 | 99.21%, 99.21% | 171,859,247,896 | 54.78X | 90.16% | 320 | 48.08 | 39.31% |
| 10138 | 1,413,986,946 | 99.36%, 99.36% | 138,460,912,489 | 44.14X | 89.31% | 323 | 48.17 | 39.73% |
| 10143 | 1,627,071,352 | 77.51%, 77.51% | 125,253,967,485 | 39.93X | 89.66% | 362 | 48.15 | 40.09% |
| 10144 | 1,964,930,229 | 68.61%, 68.61% | 133,744,891,260 | 42.63X | 89.05% | 361 | 48.21 | 40.08% |
| 10145 | 1,213,529,960 | 98.51%, 98.51% | 118,888,431,070 | 37.9X | 89.3% | 362 | 48.2 | 39.92% |
| 10231 | 1,204,598,812 | 98.12%, 98.12% | 117,678,296,174 | 37.51X | 87.54% | 331 | 48.19 | 40.06% |
| 10232 | 1,206,286,495 | 98.35%, 98.35% | 118,169,799,321 | 37.67X | 87.58% | 328 | 48.2 | 40.14% |
| 10233 | 1,215,246,598 | 98.3%, 98.3% | 118,899,663,206 | 37.9X | 87.83% | 328 | 48.2 | 40.34% |
| 10235 | 1,614,404,561 | 77.26%, 77.26% | 124,189,833,293 | 39.59X | 89.7% | 329 | 48.11 | 40.62% |

**Table S2.**
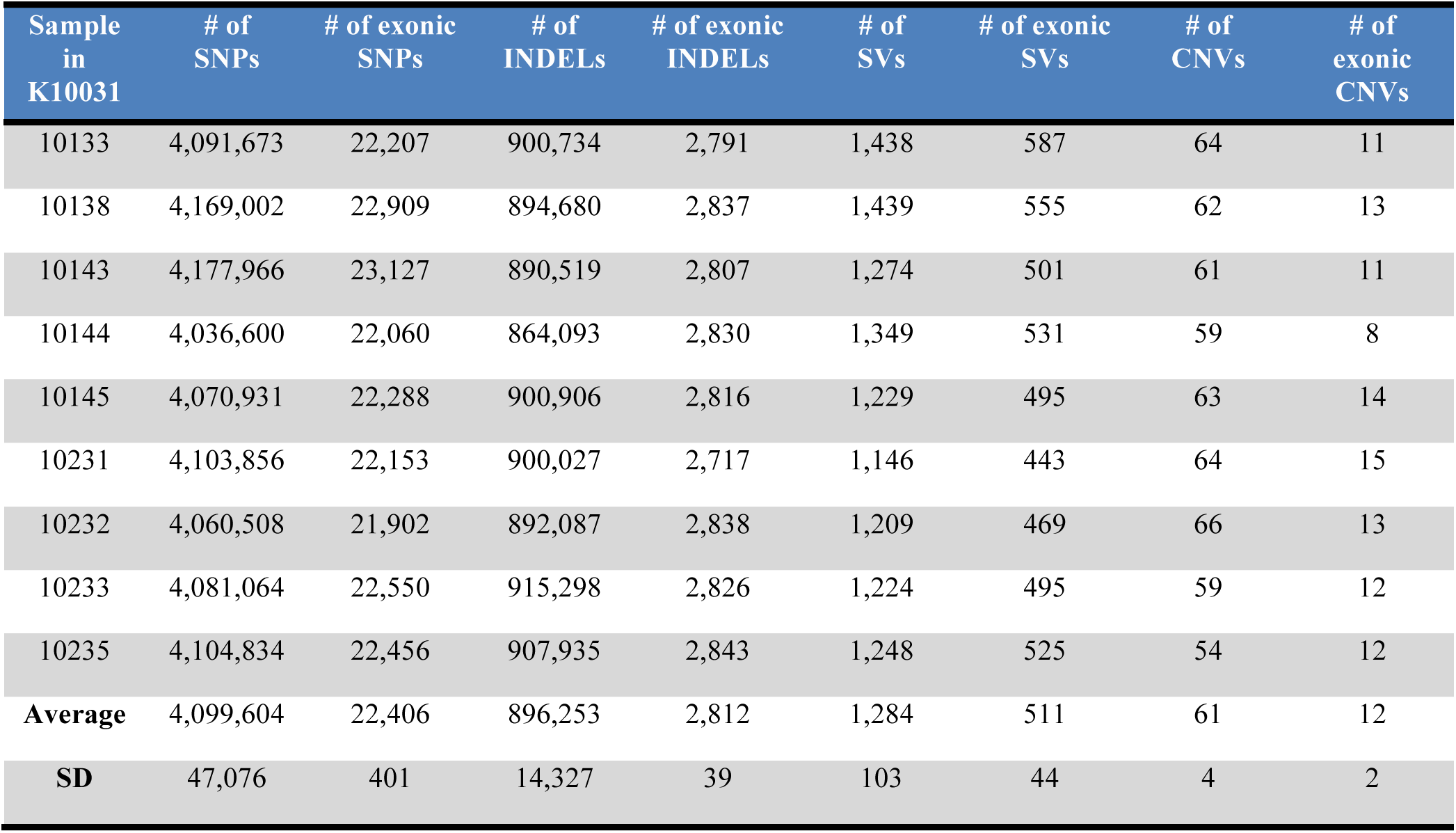
Summary statistics of the variants detected from each individual in this study.

| Sample<br>in<br>K10031 | # of<br>SNPs | # of exonic<br>SNPs | # of<br>INDELs | # of exonic<br>INDELs | # of<br>SVs | # of exonic<br>SVs | # of<br>CNVs | # of<br>exonic<br>CNVs |
| --- | --- | --- | --- | --- | --- | --- | --- | --- |
| 10133 | 4,091,673 | 22,207 | 900,734 | 2,791 | 1,438 | 587 | 64 | 11 |
| 10138 | 4,169,002 | 22,909 | 894,680 | 2,837 | 1,439 | 555 | 62 | 13 |
| 10143 | 4,177,966 | 23,127 | 890,519 | 2,807 | 1,274 | 501 | 61 | 11 |
| 10144 | 4,036,600 | 22,060 | 864,093 | 2,830 | 1,349 | 531 | 59 | 8 |
| 10145 | 4,070,931 | 22,288 | 900,906 | 2,816 | 1,229 | 495 | 63 | 14 |
| 10231 | 4,103,856 | 22,153 | 900,027 | 2,717 | 1,146 | 443 | 64 | 15 |
| 10232 | 4,060,508 | 21,902 | 892,087 | 2,838 | 1,209 | 469 | 66 | 13 |
| 10233 | 4,081,064 | 22,550 | 915,298 | 2,826 | 1,224 | 495 | 59 | 12 |
| 10235 | 4,104,834 | 22,456 | 907,935 | 2,843 | 1,248 | 525 | 54 | 12 |
| **Average** | 4,099,604 | 22,406 | 896,253 | 2,812 | 1,284 | 511 | 61 | 12 |
| **SD** | 47,076 | 401 | 14,327 | 39 | 103 | 44 | 4 | 2 |

**Table S3.** Kinship inference between each pair of the individuals using KING. A kinship score of 0.25 indicates that these two individuals are parent-child or full siblings

| FID1 | FID2 | Number_SNP | HetHet | IBS0 | Kinship |
| --- | --- | --- | --- | --- | --- |
| K10031_10144 | K10031_10145 | 7712483 | 0.155 | 0.061 | 0.0377 |
| K10031_10133 | K10031_10144 | 7720872 | 0.204 | 0.005 | 0.2459 |
| K10031_10133 | K10031_10145 | 7719024 | 0.201 | 0.0051 | 0.2478 |
| K10031_10138 | K10031_10144 | 7721400 | 0.205 | 0.0052 | 0.2478 |
| K10031_10138 | K10031_10145 | 7720609 | 0.2 | 0.0054 | 0.2458 |
| K10031_10143 | K10031_10144 | 7719412 | 0.207 | 0.0041 | 0.2527 |
| K10031_10143 | K10031_10145 | 7721142 | 0.206 | 0.0041 | 0.252 |
| K10031_10235 | K10031_10144 | 7715502 | 0.207 | 0.004 | 0.2523 |
| K10031_10235 | K10031_10145 | 7716964 | 0.201 | 0.0041 | 0.2488 |
| K10031_10231 | K10031_10145 | 7712110 | 0.224 | 0.0197 | 0.2398 |
| K10031_10231 | K10031_10232 | 7709621 | 0.202 | 0.0049 | 0.2321 |
| K10031_10231 | K10031_10233 | 7710754 | 0.203 | 0.0045 | 0.2336 |

### Supplemental data 2

**Figure S3.**
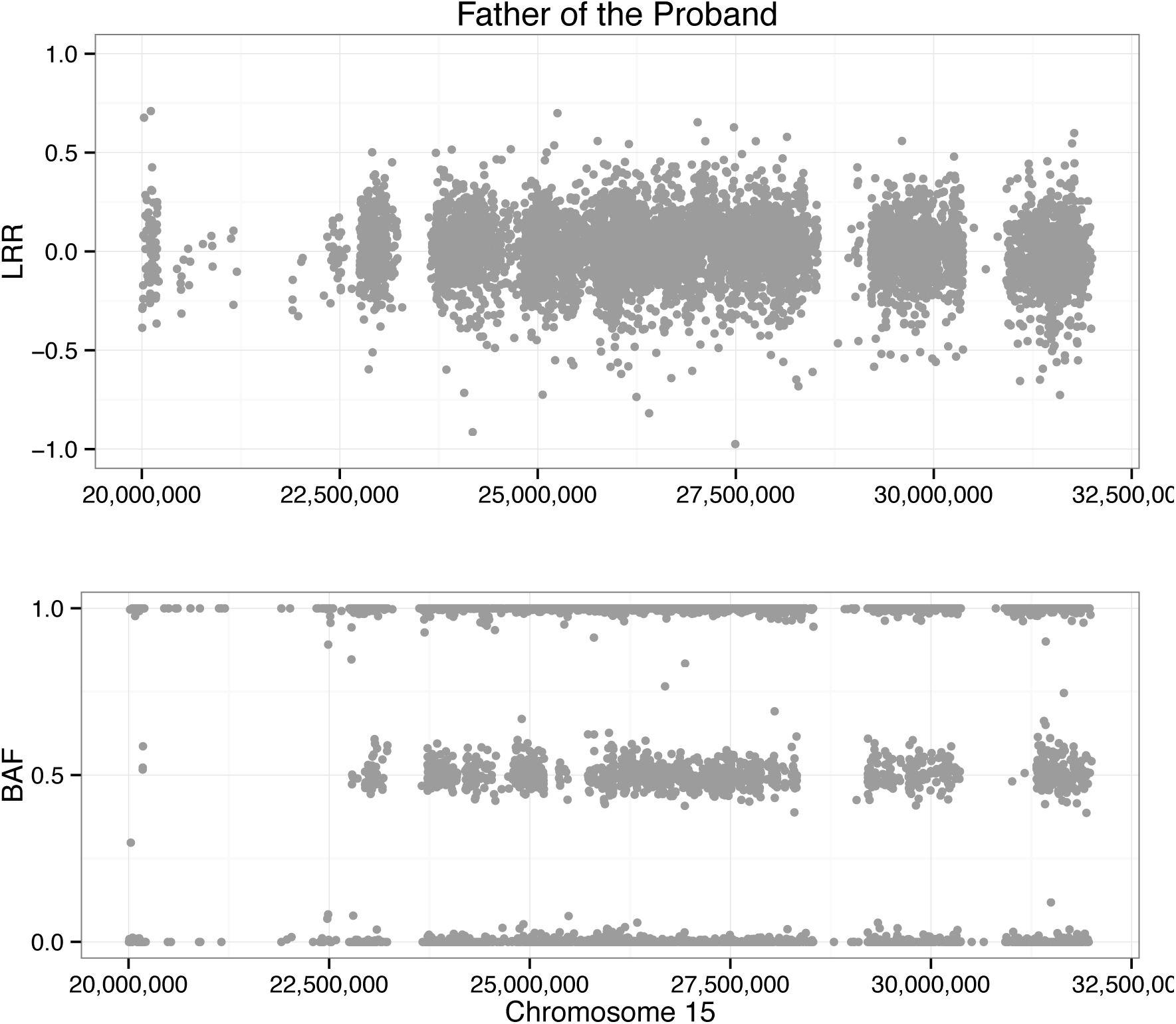
Distributions of Log-R ratios (LRR) and B allele frequencies (BAF) in 15q11.2-15q13 of the microrarray data from the father of the proband (K10031-10231) with PWS. We used PennCNV to inspect this region from Illumina 2.5m microarray data.

**Figure S4.**
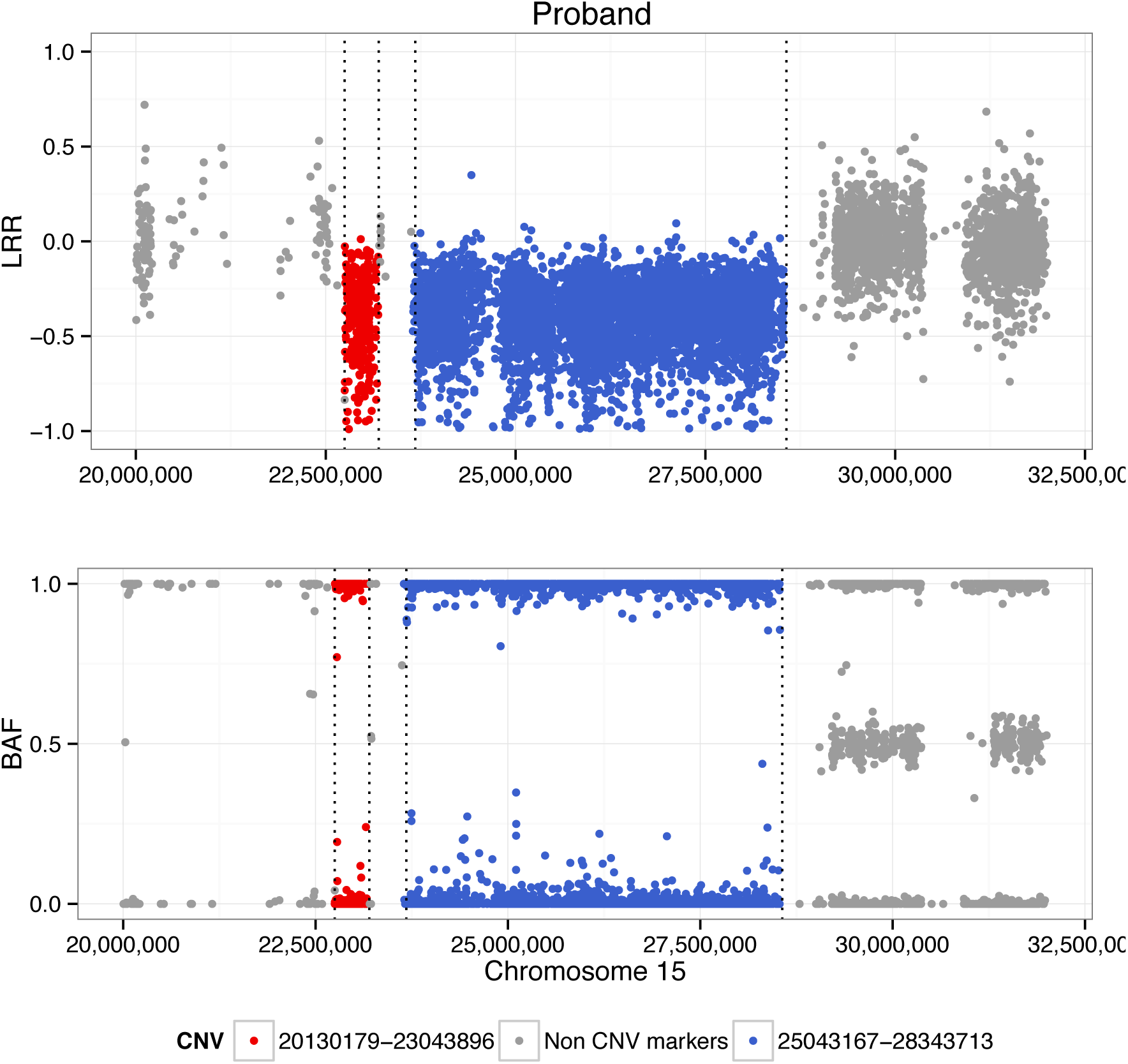
Validation of the copy number variant in the proband with PWS (K10031-10232) using Illumina 2.5m microarray data. We used PennCNV to call this deletion from the microarray data, which is also only detected from the proband, but not from the father and the two unaffected brothers. The dash lines in the figure of proband indicate the interval of the ERDS copy number variant call.

**Figure S5.**
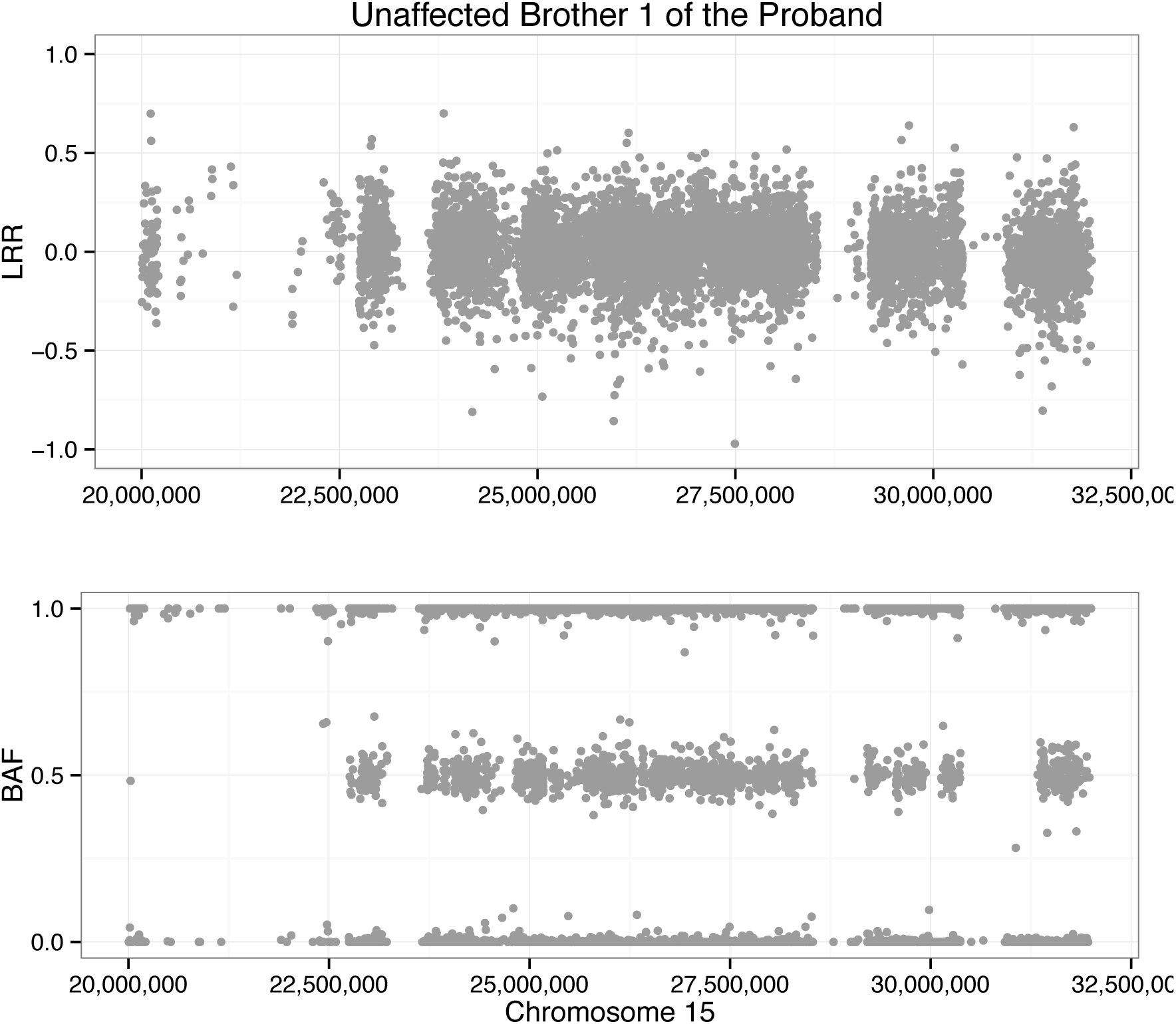
Distributions of Log-R ratios (LRR) and B allele frequencies (BAF) in 15q11.2-15q13 of the microrarray data from the unaffected brother (K10031-10233) of the proband with PWS. We used PennCNV to inspect this region from Illumina 2.5m microarray data.

**Figure S6.**
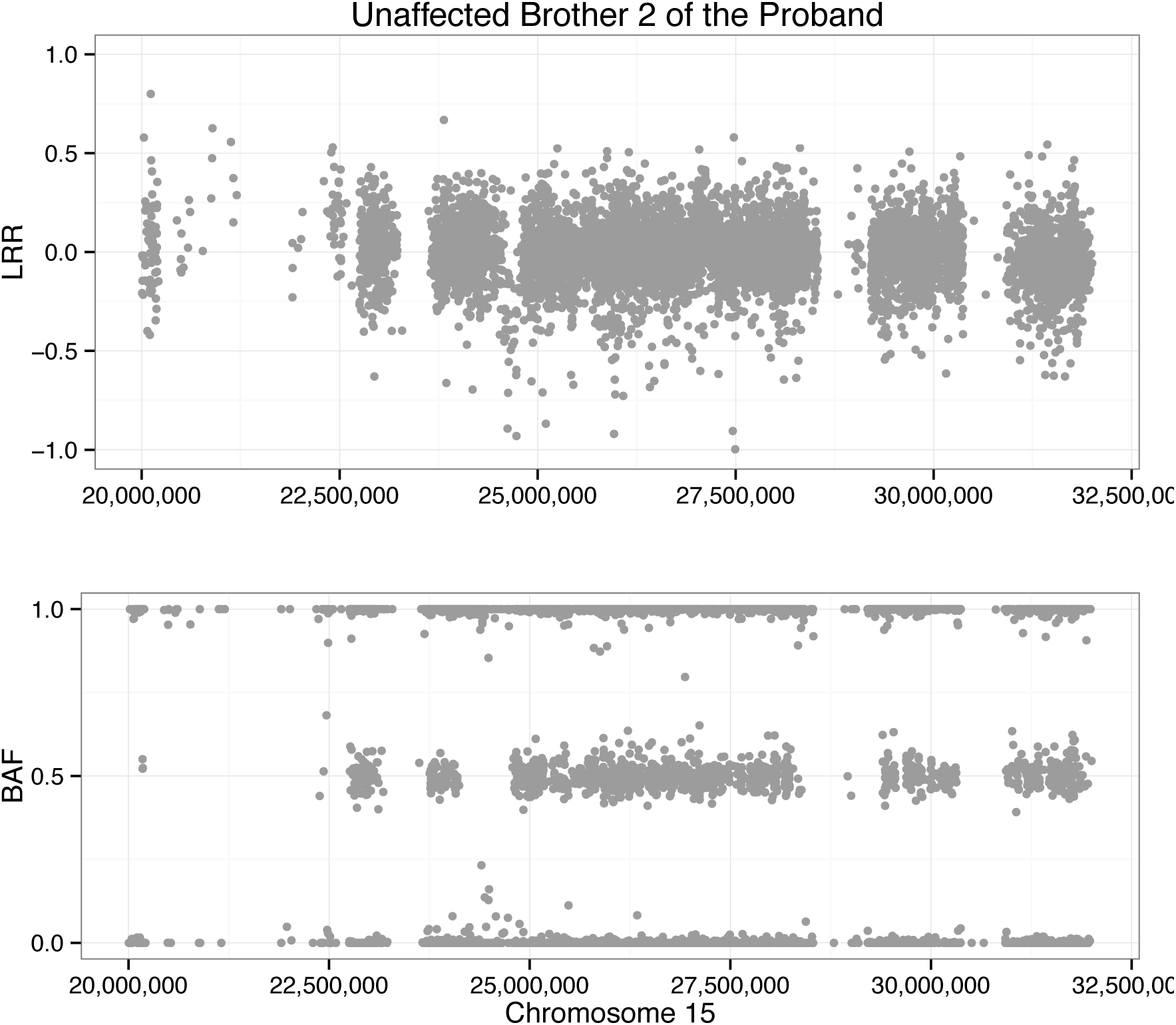
Distributions of Log-R ratios (LRR) and B allele frequencies (BAF) in 15q11.2-15q13 of the microrarray data from the unaffected brother (K10031-10234) of the proband with PWS. We used PennCNV to inspect this region from Illumina 2.5m microarray data.

**Figure S7.**
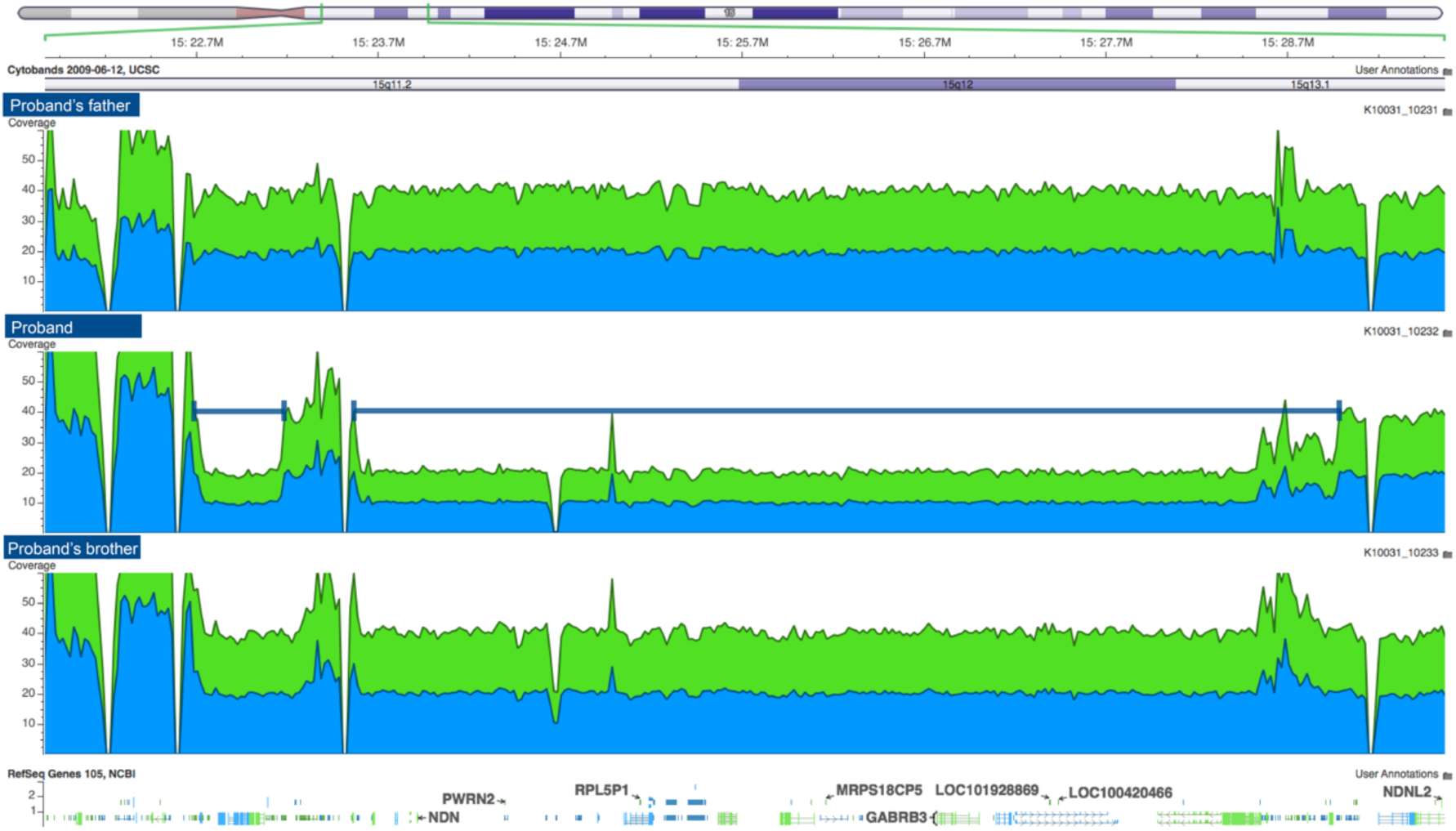
Screenshot of the alignment in 15q11.2-15q13. These deletions are not detected either in the proband’s father (K10031-10231) or the unaffected brother (K10031-10233). The deletion was confirmed with the Illumina Omni 2.5m microarray data.

### Supplemental data 3

**Table S4.** A list of variants (AAF<1% in ExAC and 1000G) found in the probands with dysautonomia-like symptoms. The coordinates are shown with respect to hg19. These variants are reported in the probands (K10031-10133, 10031-10138, K10031-10145), called by at least one pipeline, located within coding regions, are ranked high by pVAAST, and have a CADD c-score greater than 15 or have a GEMINI predicted of at least MED. *De novo* variants that are detected in K10031-10133 and/or K10031-10138 are also reported in this table, labeled with stars. The Alternative Allele Frequency (AAF) is computed based on the general population in the ExAC database or 1000G for some intronic variants.

| Gene | Genomic coordinates | Change | Effect | CADD | Zygotity & AAF |
| --- | --- | --- | --- | --- | --- |
| *ATXN2* | chr12: 112037180 | G>T | missense | 7.4 | Het, None |
| *LRRIQ1* | chr12: 85450243 | G>A | missense | 10.2 | Het, 0.37% |
| *MYO1H* | chr12: 109862622 | A>G | missense | 19.5 | Het, 0.002% |
| *OR1J4* | chr9: 125281999 | C>T | missense | 7.4 | Het, 0.62% |
| *PLCG2* | chr16: 81927314 | G>A | splice_region | 10.3 | Het, 0.1% |
| *RFX4* | chr12: 107155117 | C>T | missense | 16.67 | Het, 0.68% |
| *\*HHIPL2* | chr1: 222717014 | T>G | missense | 21.3 | Het, 0.0008% |
| *\*KIF11* | chr10: 94410261 | T>G | missense | 34.0 | Het, 0.025% |
| *\*PDZRN4* | chr12: 41883326 | C>T | intron_variant | 19.1 | Het, None |
| *\*NALCN* | chr13: 101828755 | A>C | intron_variant | 18.9 | Het, 0.007% |

### Additional Clinical Information of Individuals in the Study

#### K10031-10232

test results

#### Autism Diagnostic Observation Schedule-Generic (ADOS)-Module 2

Communication Score: 4

Social Interaction Score: 4

Stereotyped Behaviors & Restricted Interests: 0

Other Abnormal Behaviors: 4

Overall Score: 12

K10031-10232’s Language and Communication activities displayed some relatively complex language, but there were also grammatical errors, and intonation and volume varied across contexts. His conversation took place through role-playing, and some of his gestures were exaggerated, limited, or inappropriate to the given situation. For the reciprocal social interaction domain, he was able to make strong eye contact to initiate or terminate an activity with the staff, yet his facial expressions were usually out of context. He needed to be redirected on less important tasks by the examiner; when a task was broken into smaller steps and “farm animals” asked him to complete it, he engaged quickly and completed the task successfully. In the imagination domain, K10031-10232 displays spontaneity, creativity, and make-believe actions. It is important to take into consideration that K10031-10232 was markedly anxious during this assessment, which may have affected the reliability of the scores.

#### The Childhood Autism Rating Scale (CARS)

The parents of K10031-10232 yielded a score of 37.5, which falls in the high end of mildly-moderately autistic. The parents noted that their son’s intellectual level is abnormal in that he is not as intelligent as a “normally developing child”, but he may function better than the “normal child” of the same age in one or more areas. The parents also rated his emotional response to specific situations as inappropriate and inhibited (score of 3 for Emotional Responses). They also noted intense and frequent abnormal body movements (3.5 score for Body Use).

#### K10031-10138

Individual K10031-10138, K10031-10133’s brother, also presents with dysautonomia-like symptoms. He is put in a wheelchair to prevent the injuries from frequent syncopal events. He has bradycardia and tachycardia. Individual K10031-10138 is also diagnosed with Tourette Syndrome; observable tics include tapping, abdominal crunches, vocal sniffing and grunting, and eyeblinking. And his ADD makes it difficult to focus in school. Moreover, he has OCD too; everything must be clean and “just right”, and needs a private bathroom.

#### K10031-10261

Individual K10031-10261, K10031-10133’s another brother, has dark brown hair, pale skin, is overweight. He has asthma, seizures in response to Pertussis, dyslexia, migraines, arthritis, is very forgetful, has ADD but is not hyper, and has had two syncopic episodes. He is able to calmly give accounts of the two times he passed out, and describes both experiences as dizzying, losing consciousness, and waking up on the floor. See HDV_0072 for a clinical presentation of K10031-10261.

#### K10031-10145

Individual K10031-10145, the mother of the proband K10031-10133, has hemochromatosis and dysautonomia-like symptomes, which is claimed to have been inherited from her biological mother. Individual K10031-10145’s dysautonomia-like symptoms started during her anorexia issues at age 19. She experiences dizziness frequently and has OCD (is always checking the curling irons to ensure they are turned off). She is also diagnosed with anxiety, depression, osteoporosis, dissociative identity disorder (DID) which makes her moody and resulted in a period of suicidal thoughts.

#### K10031-10143

Individual K10031-10143, K10031-10133’s sister, is quite normal other than her separation anxiety. She does get dizzy but only passed out once.

#### K10031-10235

Individual K10031-10235, K10031-10133’s sister, has acid reflux, tremors, migraines, asthma, anxiety, and possibly bipolar disorder. The bipolar disorder is suspected due to family members’ concerns for her mood swings, spending sprees, and irritability. See description of supplemental videos HDV_0080 and HDV_0081 at the end of document for further details. These videos can be made available upon request to qualified investigators.

#### K10031-10144

Individual K10031-10144, K10031-10133’s father, has migraines, arthritis, gastroesophageal reflux disease (GERD), and hiatal hernia. He has ADD, a history of suicidal attempt, and has two sisters with mental illnesses.

#### K10031-10233

Individual K10031-10233, first cousin to K10031-10133 and brother to K10031-10232, is diagnosed with ADHD and depression. It is worth noting that K10031-10233’s other brother, K10031-10234, is also diagnosed with ADD, as well as bipolar disorder and Tourette syndrome.

### Supplemental Videos (Description of Content. Videos can be made available upon request to qualified investigators)

#### HDV_0079

Video Summary: In this video, the proband (K10031-10133) introduces her current medical conditions, including dysautonomia-like symptoms, bradycardia, tachycardia, heat intolerance, gastrophoresis, stroke, PFO, and glaucoma. She denies any psychological diagnosis on herself such as OCD and Tourette syndrome. Her parents (K10031-10145/10231) also discussed their family history of osteoporosis, Tourette’s and bipolar disorder.

#### HDV_0072

Video Summary: In this video, more relatives of the proband (K10031-10133) were introduced, including her mother (K10031-10145), her maternal uncle (K10031-10231), her younger sisters (K10031-10235/10143), her younger brother (K10031-10261), and her cousin (K10031-10232). Medical history was collected from her younger brother (10261), who reports to have slight blood pressure issues, two fainting episodes and ADD, one of her younger sister (10235) and her cousin with Prader-Willi syndrome (PWS) (K10031-10232).

#### HDV_0073

Video Summary: More interviews on other proband with PWS (K10031-10232) and his father (K10031-10231) were made in this video. His father (10231) explains the medications his son is taking, including growth hormone, Abilify, and Sertraline, and certain types of behaviors his son presents before taking his medications. Also, the mother of the female proband (K10031-10133) explains more about her own medical diagnoses, including hemochromatosis and low blood pressure, as well as her children’s issues with dysautonomia-like symptoms. Another maternal uncle (K10031-10262) was also interviewed in this video, who explains the negative medical and family histories of his own family. During the interview, his verbal consent of participation in this study was also collected.

## REFERENCES

Abyzov A, Urban AE, Snyder M, Gerstein M. 2011. CNVnator: An approach to discover, genotype, and characterize typical and atypical CNVs from family and population genome sequencing. Genome Research 21: 974–974.

Allen KJ, Gurrin LC, Constantine CC, Osborne NJ, Delatycki MB, Nicoll AJ, McLaren CE, Bahlo M, Nisselle AE, Vulpe CD et al. 2008. Iron-Overload–Related Disease in HFE Hereditary Hemochromatosis. New England Journal of Medicine 358: 221–221.

Bamshad MJ, Ng SB, Bigham AW, Tabor HK, Emond MJ, Nickerson DA, Shendure J. 2011. Exome sequencing as a tool for Mendelian disease gene discovery. Nat Rev Genet 12: 745–745.

Beutler E, Felitti VJ, Koziol JA, Ho NJ, Gelbart T. 2002. Penetrance of 845G→A (C282Y) HFE hereditary haemochromatosis mutation in the USA. The Lancet 359: 211–211.

Bieth E, Eddiry S, Gaston V, Lorenzini F, Buffet A, Conte Auriol F, Molinas C, Cailley D, Rooryck C, Arveiler B et al. 2014. Highly restricted deletion of the SNORD116 region is implicated in Prader-Willi Syndrome. Eur J Hum Genet doi:10.1038/ejhg.2014.103.

Boycott KM, Vanstone MR, Bulman DE, MacKenzie AE. 2013. Rare-disease genetics in the era of next-generation sequencing: discovery to translation. Nat Rev Genet 14: 681–681.

Cantor SB, Bell DW, Ganesan S, Kass EM, Drapkin R, Grossman S, Wahrer DCR, Sgroi DC, Lane WS, Haber DA et al. 2001. BACH1, a Novel Helicase-like Protein, Interacts Directly with BRCA1 and Contributes to Its DNA Repair Function. Cell 105: 149–160.

Cargill M, Altshuler D, Ireland J, Sklar P, Ardlie K, Patil N, Lane CR, Lim EP, Kalyanaraman N, Nemesh J et al. 1999. Characterization of single-nucleotide polymorphisms in coding regions of human genes. Nat Genet 22: 231–231.

Cassidy SB, Driscoll DJ. 2008. Prader-Willi syndrome. Eur J Hum Genet 17: 3–13.

Cassidy SB, Schwartz S, Miller JL, Driscoll DJ. 2012. Prader-Willi syndrome. Genet Med 14: 10–10.

Christenhusz GM, Devriendt K, Dierickx K. 2013. Secondary variants - in defense of a more fitting term in the incidental findings debate. Eur J Hum Genet 21:1331–1334.

Christian SL, Robinson WP, Huang B, Mutirangura A, Line MR, Nakao M, Surti U, Chakravarti A, Ledbetter DH. 1995. Molecular characterization of two proximal deletion breakpoint regions in both Prader-Willi and Angelman syndrome patients. American Journal of Human Genetics 57: 40–40.

Corbett-Detig RB, Zhou J, Clark AG, Hartl DL, Ayroles JF. 2013. Genetic incompatibilities are widespread within species. Nature 504: 135–135.

DePristo MA, Banks E, Poplin R, Garimella KV, Maguire JR, Hartl C, Philippakis AA, del Angel G, Rivas MA, Hanna M et al. 2011. A framework for variation discovery and genotyping using next-generation DNA sequencing data. Nat Genet 43: 491–491.

Dewey FE, Grove ME, Pan C, et al. 2014. CLinical interpretation and implications of whole-genome sequencing. JAMA 311: 1035–1035.

Dittrich B, Robinson W, Knoblauch H, Buiting K, Schmidt K, Gillessen-Kaesbach G, Horsthemke B. 1992. Molecular diagnosis of the Prader-Willi and Angelman syndromes by detection of parent-of-origin specific DNA methylation in 15q11-13. Hum Genet 90: 313–313.

Esteller M. 2011. Non-coding RNAs in human disease. Nat Rev Genet 12: 861–874.

Evans JP, Skrzynia C, Burke W. 2001. The complexities of predictive genetic testing. BMJ : British Medical Journal 322: 1052–1052.

Fang H, Wu Y, Narzisi G, O’Rawe J, Barrón LTJ, Rosenbaum J, Ronemus M, Iossifov I, Schatz M, Lyon G. 2014. Reducing INDEL calling errors in whole-genome and exome sequencing data. Genome Medicine: 89.

Faustino NA, Cooper TA. 2003. Pre-mRNA splicing and human disease. Genes & Development 17: 419–419.

Freund J, Brandmaier AM, Lewejohann L, Kirste I, Kritzler M, Krüger A, Sachser N, Lindenberger U, Kempermann G. 2013. Emergence of Individuality in Genetically Identical Mice. Science 340: 756–756.

G. Day-Williams A, Sun C, Jelcic I, McLaughlin H, Harris T, Martin R, Carulli J. 2015. Whole Genome Sequencing Reveals a Chromosome 9p Deletion Causing DOCK8 Deficiency in an Adult Diagnosed with Hyper IgE Syndrome Who Developed Progressive Multifocal Leukoencephalopathy. J Clin Immunol 35: 92–92.

García-Alcalde F, Okonechnikov K, Carbonell J, Cruz LM, Götz S, Tarazona S, Dopazo J, Meyer TF, Conesa A. 2012. Qualimap: evaluating next-generation sequencing alignment data. Bioinformatics 28: 2678–2678.

Garrison E, Marth G. 2012. Haplotype-based variant detection from short-read sequencing. ArXiv e-prints 1207: 3907.

Gimm O, Greco A, Hoang-Vu C, Dralle H, Pierotti MA, Eng C. 1999. Mutation Analysis Reveals Novel Sequence Variants in NTRK1 in Sporadic Human Medullary Thyroid Carcinoma. The Journal of Clinical Endocrinology & Metabolism 84: 2784–2784.

Greenman C, Stephens P, Smith R, Dalgliesh GL, Hunter C, Bignell G, Davies H, Teague J, Butler A, Stevens C et al. 2007. Patterns of somatic mutation in human cancer genomes. Nature 446: 153–153.

Grillo E, Lo Rizzo C, Bianciardi L, Bizzarri V, Baldassarri M, Spiga O, Furini S, De Felice C, Signorini C, Leoncini S et al. 2013. Revealing the complexity of a monogenic disease: rett syndrome exome sequencing. PLoS One 8: e56599.

Gu Z, Gu L, Eils R, Schlesner M, Brors B. 2014. circlize implements and enhances circular visualization in R. Bioinformatics 30: 2811–2811.

Hamilton BA, Yu BD. 2012. Modifier genes and the plasticity of genetic networks in mice. PLoS genetics 8: e1002644.

Hanson EH, Imperatore G, Burke W. 2001. HFE Gene and Hereditary Hemochromatosis: A HuGE Review. American Journal of Epidemiology 154: 193–193.

Hewett M, Oliver DE, Rubin DL, Easton KL, Stuart JM, Altman RB, Klein TE. 2002. PharmGKB: the Pharmacogenetics Knowledge Base. Nucleic Acids Research 30: 163–163.

Highnam G, Franck C, Martin A, Stephens C, Puthige A, Mittelman D. 2013. Accurate human microsatellite genotypes from high-throughput resequencing data using informed error profiles. Nucleic Acids Research 41: e32.

Honeyman JN, Simon EP, Robine N, Chiaroni-Clarke R, Darcy DG, Lim IIP, Gleason CE, Murphy JM, Rosenberg BR, Teegan L et al. 2014. Detection of a Recurrent DNAJB1-PRKACA Chimeric Transcript in Fibrolamellar Hepatocellular Carcinoma. Science 343: 1010–1010.

Hu H, Roach JC, Coon H, Guthery SL, Voelkerding KV, Margraf RL, Durtschi JD, Tavtigian SV, Shankaracharya, Wu W et al. 2014. A unified test of linkage analysis and rare-variant association for analysis of pedigree sequence data. Nat Biotech 32: 663–663.

Indo Y. 2001. Molecular basis of congenital insensitivity to pain with anhidrosis (CIPA): Mutations and polymorphisms in TRKA (NTRK1) gene encoding the receptor tyrosine kinase for nerve growth factor. Human Mutation 18: 462–462.

Indo Y, Tsuruta M, Hayashida Y, Karim MA, Ohta K, Kawano T, Mitsubuchi H, Tonoki H, Awaya Y, Matsuda I. 1996. Mutations in the TRKA/NGF receptor gene in patients with congenital insensitivity to pain with anhidrosis. Nat Genet 13: 485–485.

Johnson JA, Gong L, Whirl-Carrillo M, Gage BF, Scott SA, Stein CM, Anderson JL, Kimmel SE, Lee MTM, Pirmohamed M et al. 2011. Clinical Pharmacogenetics Implementation Consortium Guidelines for CYP2C9 and VKORC1 Genotypes and Warfarin Dosing. Clin Pharmacol Ther 90: 625–625.

Kircher M, Witten DM, Jain P, O’Roak BJ, Cooper GM, Shendure J. 2014. A general framework for estimating the relative pathogenicity of human genetic variants. Nat Genet 46: 310–310.

Knoll JHM, Nicholls RD, Magenis RE, Graham JM, Lalande M, Latt SA, Opitz JM, Reynolds JF. 1989. Angelman and Prader-Willi syndromes share a common chromosome 15 deletion but differ in parental origin of the deletion. American Journal of Medical Genetics 32: 285–285.

Koboldt Daniel C, Steinberg Karyn M, Larson David E, Wilson Richard K, Mardis ER. 2013. The Next-Generation Sequencing Revolution and Its Impact on Genomics. Cell 155: 27–27.

Kohler S, Doelken SC, Mungall CJ, Bauer S, Firth HV, Bailleul-Forestier I, Black GC, Brown DL, Brudno M, Campbell J et al. 2014. The Human Phenotype Ontology project: linking molecular biology and disease through phenotype data. Nucleic Acids Res 42: D966–974.

Köhler S, Schoeneberg U, Czeschik JC, Doelken SC, Hehir-Kwa JY, Ibn-Salem J, Mungall CJ, Smedley D, Haendel MA, Robinson PN. 2014. Clinical interpretation of CNVs with cross-species phenotype data. Journal of Medical Genetics doi:10.1136/jmedgenet-2014–102633.

Kohler S, Schulz MH, Krawitz P, Bauer S, Dolken S, Ott CE, Mundlos C, Horn D, Mundlos S, Robinson PN. 2009. Clinical diagnostics in human genetics with semantic similarity searches in ontologies. Am J Hum Genet 85: 457–457.

Layer R, Chiang C, Quinlan A, Hall I. 2014. LUMPY: a probabilistic framework for structural variant discovery. Genome Biology 15: R84.

Lee H, Deignan JL, Dorrani N, et al. 2014. CLinical exome sequencing for genetic identification of rare mendelian disorders. JAMA 312: 1880–1880.

Li H. 2013. Aligning sequence reads, clone sequences and assembly contigs with BWA-MEM. ArXiv e-prints 1303: 3997.

Li H, Cherry S, Klinedinst D, DeLeon V, Redig J, Reshey B, Chin MT, Sherman SL, Maslen CL, Reeves RH. 2012. Genetic modifiers predisposing to congenital heart disease in the sensitized Down syndrome population. Circ Cardiovasc Genet 5: 301–301.

Li H, Handsaker B, Wysoker A, Fennell T, Ruan J, Homer N, Marth G, Abecasis G, Durbin R, Subgroup GPDP. 2009. The Sequence Alignment/Map format and SAMtools. Bioinformatics 25: 2078–2078.

Lyon G, Wang K. 2012. Identifying disease mutations in genomic medicine settings: current challenges and how to accelerate progress. Genome Medicine 4: 58.

Lyon GJ. 2012a. Personalized medicine: Bring clinical standards to human-genetics research. Nature 482: 300–300.

Lyon GJ. 2012b. There is nothing ‘incidental’ about unrelated findings. Personalized Medicine 9: 163–163.

Lyon GJ, O’Rawe J. 2014. Human genetics and clinical aspects of neurodevelopmental disorders. In The Genetics of Neurodevelopmental Disorders, doi:10.1101/000687 (ed. K Mitchell). Cold Spring Harbor Labs Journals.

Lyon GJ, Segal JP. 2013. Practical, ethical and regulatory considerations for the evolving medical and research genomics landscape. Applied & Translational Genomics 2: 34–34.

MacArthur DG, Balasubramanian S, Frankish A, Huang N, Morris J, Walter K, Jostins L, Habegger L, Pickrell JK, Montgomery SB et al. 2012. A Systematic Survey of Loss-of-Function Variants in Human Protein-Coding Genes. Science 335: 823–823.

Mackay TFC, Stone EA, Ayroles JF. 2009. The genetics of quantitative traits: challenges and prospects. Nat Rev Genet 10: 565–565.

Malcolm S, Clayton-Smith J, Nichols M, Pembrey ME, Armour JAL, Jeffreys AJ, Robb S, Webb T. 1991. Uniparental paternal disomy in Angelman’s syndrome. The Lancet 337: 694–694.

Manichaikul A, Mychaleckyj JC, Rich SS, Daly K, Sale M, Chen W-M. 2010. Robust relationship inference in genome-wide association studies. Bioinformatics 26: 2867–2867.

Mardy S, Miura Y, Endo F, Matsuda I, Indo Y. 2001. Congenital insensitivity to pain with anhidrosis (CIPA): effect of TRKA (NTRK1) missense mutations on autophosphorylation of the receptor tyrosine kinase for nerve growth factor. Human Molecular Genetics 10: 179–179.

Massouras A, Waszak SM, Albarca-Aguilera M, Hens K, Holcombe W, Ayroles JF, Dermitzakis ET, Stone EA, Jensen JD, Mackay TFC et al. 2012. Genomic Variation and Its Impact on Gene Expression in Drosophila melanogaster. PLoS Genet 8: e1003055.

McKenna A, Hanna M, Banks E, Sivachenko A, Cibulskis K, Kernytsky A, Garimella K, Altshuler D, Gabriel S, Daly M et al. 2010. The Genome Analysis Toolkit: A MapReduce framework for analyzing next-generation DNA sequencing data. Genome Research 20: 1297–1297.

Meijers-Heijboer EJ, Verhoog LC, Brekelmans CTM, Seynaeve C, Tilanus-Linthorst MMA, Wagner A, Dukel L, Devilee P, van den Ouweland AMW, van Geel AN et al. 2000. Presymptomatic DNA testing and prophylactic surgery in families with a BRCA1 or BRCA2 mutation. The Lancet 355: 2015–2015.

Moczulski DK, Grzeszczak W, Gawlik B. 2001. Role of Hemochromatosis C282Y and H63D Mutations in HFE Gene in Development of Type 2 Diabetes and Diabetic Nephropathy. Diabetes Care 24: 1187–1187.

Morton CC, Nance WE. 2006. Newborn Hearing Screening — A Silent Revolution. New England Journal of Medicine 354: 2151–2151.

Nanda R, Schumm L, Cummings S, et al. 2005. Genetic testing in an ethnically diverse cohort of high-risk women: A comparative analysis of brca1 and brca2 mutations in american families of european and african ancestry. JAMA 294: 1925–1925.

Narzisi G, O’Rawe JA, Iossifov I, Fang H, Lee Y-h, Wang Z, Wu Y, Lyon GJ, Wigler M, Schatz MC. 2014. Accurate de novo and transmitted indel detection in exome-capture data using microassembly. Nat Meth advance online publication.

Nicholls RD, Knepper JL. 2001. GENOME ORGANIZATION, FUNCTION, AND IMPRINTING IN PRADER-WILLI AND ANGELMAN SYNDROMES. Annual Review of Genomics and Human Genetics 2: 153–153.

O’Rawe J, Jiang T, Sun G, Wu Y, Wang W, Hu J, Bodily P, Tian L, Hakonarson H, Johnson WE et al. 2013a. Low concordance of multiple variant-calling pipelines: practical implications for exome and genome sequencing. Genome Medicine 5: 28.

O’Rawe JA, Fang H, Rynearson S, Robison R, Kiruluta ES, Higgins G, Eilbeck K, Reese MG, Lyon GJ. 2013d. Integrating precision medicine in the study and clinical treatment of a severely mentally ill person. PeerJ 1: e177.

Pagani F, Stuani C, Tzetis M, Kanavakis E, Efthymiadou A, Doudounakis S, Casals T, Baralle FE. 2003. New type of disease causing mutations: the example of the composite exonic regulatory elements of splicing in CFTR exon 12. Human Molecular Genetics 12: 1111–1111.

Paila U, Chapman BA, Kirchner R, Quinlan AR. 2013. GEMINI: Integrative Exploration of Genetic Variation and Genome Annotations. PLoS Comput Biol 9: e1003153.

Palomaki GE, Kloza EM, Lambert-Messerlian GM, Haddow JE, Neveux LM, Ehrich M, van den Boom D, Bombard AT, Deciu C, Grody WW et al. 2011. DNA sequencing of maternal plasma to detect Down syndrome: An international clinical validation study. Genet Med 13: 913–913.

Pietrangelo A. 2004. Hereditary Hemochromatosis — A New Look at an Old Disease. New England Journal of Medicine 350: 2383–2383.

Quinlan AR, Hall IM. 2010. BEDTools: a flexible suite of utilities for comparing genomic features. Bioinformatics 26: 841–841.

Robinson PN, Köhler S, Bauer S, Seelow D, Horn D, Mundlos S. 2008. The Human Phenotype Ontology: A Tool for Annotating and Analyzing Human Hereditary Disease. The American Journal of Human Genetics 83: 610–610.

Robinson PN, Mundlos S. 2010. The Human Phenotype Ontology. Clinical Genetics 77: 525–525.

Rope Alan F, Wang K, Evjenth R, Xing J, Johnston Jennifer J, Swensen Jeffrey J, Johnson WE, Moore B, Huff Chad D, Bird Lynne M et al. 2011. Using VAAST to Identify an X-Linked Disorder Resulting in Lethality in Male Infants Due to N-Terminal Acetyltransferase Deficiency. The American Journal of Human Genetics 89: 28–28.

Schaaf CP, Gonzalez-Garay ML, Xia F, Potocki L, Gripp KW, Zhang B, Peters BA, McElwain MA, Drmanac R, Beaudet AL et al. 2013. Truncating mutations of MAGEL2 cause Prader-Willi phenotypes and autism. Nat Genet 45: 1405–1405.

Shatzky S, Moses S, Levy J, Pinsk V, Hershkovitz E, Herzog L, Shorer Z, Luder A, Parvari R. 2000. Congenital insensitivity to pain with anhidrosis (CIPA) in Israeli-Bedouins: Genetic heterogeneity, novel mutations in the TRKA/NGF receptor gene, clinical findings, and results of nerve conduction studies. American Journal of Medical Genetics 92: 353–353.

Sherman S, Pletcher BA, Driscoll DA. 2005. Fragile X syndrome: Diagnostic and carrier testing. Genet Med 7: 584–584.

Slaugenhaupt SA, Blumenfeld A, Gill SP, Leyne M, Mull J, Cuajungco MP, Liebert CB, Chadwick B, Idelson M, Reznik L et al. 2001. Tissue-Specific Expression of a Splicing Mutation in the IKBKAP Gene Causes Familial Dysautonomia. The American Journal of Human Genetics 68: 598–-605.

Smith GD, Ebrahim S, Lewis S, Hansell AL, Palmer LJ, Burton PR. 2005. Genetic epidemiology and public health: hope, hype, and future prospects. The Lancet 366: 1484–1484.

Stelzer Y, Sagi I, Yanuka O, Eiges R, Benvenisty N. 2014. The noncoding RNA IPW regulates the imprinted DLK1-DIO3 locus in an induced pluripotent stem cell model of Prader-Willi syndrome. Nat Genet 46: 551–551.

Stone DL, Slavotinek A, Bouffard GG, Banerjee-Basu S, Baxevanis AD, Barr M, Biesecker LG. 2000. Mutation of a gene encoding a putative chaperonin causes McKusick-Kaufman syndrome. Nat Genet 25: 79–79.

Swanson AG. 1963. Congenital insensitivity to pain with anhydrosis: A unique syndrome in two male siblings. Archives of Neurology 8: 299–299.

Teich N, Nemoda Z, Köhler H, Heinritz W, Mössner J, Keim V, Sahin-Tóth M. 2005. Gene conversion between functional trypsinogen genes PRSS1 and PRSS2 associated with chronic pancreatitis in a six-year-old girl. Human mutation 25: 343–343.

The International Warfarin Pharmacogenetics C. 2009. Estimation of the Warfarin Dose with Clinical and Pharmacogenetic Data. The New England journal of medicine 360: 753–753.

Thompson DC, McPhillips H, Davis RL, Lieu TA, Homer CJ, Helfand M. 2001. Universal newborn hearing screening: Summary of evidence. JAMA 286: 2000–2000.

Thornton-Wells TA, Moore JH, Haines JL. 2004. Genetics, statistics and human disease: analytical retooling for complexity. Trends in Genetics 20: 640–640.

Venables JP. 2004. Aberrant and Alternative Splicing in Cancer. Cancer Research 64: 7647–7647.

Walker FO. 2007. Huntington’s disease. The Lancet 369: 218–218.

Wang G-S, Cooper TA. 2007. Splicing in disease: disruption of the splicing code and the decoding machinery. Nat Rev Genet 8: 749–749.

Wang K, Kim C, Bradfield J, Guo Y, Toskala E, Otieno F, Hou C, Thomas K, Cardinale C, Lyon G et al. 2013. Whole-genome DNA/RNA sequencing identifies truncating mutations in RBCK1 in a novel Mendelian disease with neuromuscular and cardiac involvement. Genome Medicine 5: 67.

Wang K, Li M, Hadley D, Liu R, Glessner J, Grant SFA, Hakonarson H, Bucan M. 2007. PennCNV: An integrated hidden Markov model designed for high-resolution copy number variation detection in whole-genome SNP genotyping data. Genome Research 17: 1665–1665.

Wang K, Li M, Hakonarson H. 2010. ANNOVAR: functional annotation of genetic variants from high-throughput sequencing data. Nucleic Acids Research 38: e164.

Weischenfeldt J, Symmons O, Spitz F, Korbel JO. 2013. Phenotypic impact of genomic structural variation: insights from and for human disease. Nat Rev Genet 14: 125–125.

Wilke RA, Ramsey LB, Johnson SG, Maxwell WD, McLeod HL, Voora D, Krauss RM, Roden DM, Feng Q, Cooper-DeHoff RM et al. 2012. The Clinical Pharmacogenomics Implementation Consortium: CPIC Guideline for SLCO1B1 and Simvastatin-Induced Myopathy. Clin Pharmacol Ther 92: 112–112.

Yin Q-F, Yang L, Zhang Y, Xiang J-F, Wu Y-W, Carmichael Gordon G, Chen L-L. 2012. Long Noncoding RNAs with snoRNA Ends. Molecular Cell 48: 219–219.

Zemojtel T, Köhler S, Mackenroth L, Jäger M, Hecht J, Krawitz P, Graul-Neumann L, Doelken S, Ehmke N, Spielmann M et al. 2014. Effective diagnosis of genetic disease by computational phenotype analysis of the disease-associated genome. Science Translational Medicine 6: 252ra123.

Zhu M, Need AC, Han Y, Ge D, Maia JM, Zhu Q, Heinzen EL, Cirulli ET, Pelak K, He M et al. 2012. Using ERDS to infer copy-number variants in high-coverage genomes. Am J Hum Genet 91: 408–408.

Zuk O, Hechter E, Sunyaev SR, Lander ES. 2012. The mystery of missing heritability: Genetic interactions create phantom heritability. Proc Natl Acad Sci U S A 109: 1193–1193.

